# White matter microstructure and its relation to longitudinal measures of depressive symptoms in mid-late life

**DOI:** 10.1101/617530

**Authors:** Xueyi Shen, Mark J Adams, Tuula E Ritakari, Simon R Cox, Andrew M McIntosh, Heather C Whalley

## Abstract

**Background:** Studies of white matter microstructure in depression typically show alterations in depressed individuals, but they are frequently limited by small sample sizes and the absence of longitudinal measures of depressive symptoms. Depressive symptoms are however dynamic, and understanding the neurobiology of different trajectories could have important clinical implications.

**Methods:** We examined associations between current and longitudinal measures of depressive symptoms and white matter microstructure (Fractional Anisotropy, FA; Mean Diffusivity; MD) in the UK Biobank Imaging study. Depressive symptoms were assessed on 2-4 occasions over 5.9 to 10.7 years (on N=18,959 individuals on at least two occasions, N=4,444 on four occasions) from which we derived four measures of depressive symptomatology; (i) cross-sectional measure at the time of scan (imaging was conducted at a single time point), and three longitudinal measures, (ii) trajectory (iii) mean and (iv) intra-subject variance over time.

**Results:** Decreased white matter microstructure in the anterior thalamic radiation demonstrated significant associations across all four measures of depressive symptoms (for MD: β=0.020 to 0.029, p_corr_<0.030). The greatest effect sizes were however seen between decreasing white matter integrity and increasing longitudinal progression of symptoms (for MD: β=0.030 to 0.040, p_corr_<0.049). Cross-sectional symptom severity was particularly associated with decreased white matter integrity in association fibres and thalamic radiations (MD: β=0.015 to 0.039, p_corr_<0.041). While greater mean and within subject variance of depressive symptoms were mainly associated with decreased white matter microstructure within projection fibres (MD: β=0.019 to 0.029, p_corr_<0.044).

**Conclusions:** These findings indicate shared and differential neurobiological associations with severity, course and intra-subject variability of depressive symptoms. This enriches our understanding of the neurobiology underlying dynamic features of the disorder.

## Introduction

Major depressive disorder (MDD) is a disabling disorder with a heritability of approximately 37% (1; 2) and 16% lifetime risk (3). It is a heterogeneous illness(4; 5), often studied in modest sample sizes(6). These limitations have led to inconsistent findings(7) and to an uncertain relationship between quantitative measures of depressive symptoms and associated neurobiology.

A possible contributor to heterogeneous imaging findings in MDD is the longitudinal variability of depressive symptoms(8). This dynamic property is rarely captured by most imaging investigations, but potentially has important implications both in terms of understanding disease heterogeneity as well as having clinical relevance(9–11). Comparing the neurobiological associations of current and longitudinal depressive symptoms is important for identifying causal mechanisms underlying depressive symptoms, as well as identifying predictors of symptom onset, variability and progression over time. Brain structural measures have previously been found to be associated with stable depressive conditions over time, such as self-declared life-time depression(12–14). Studies on symptomatic changes however have also suggested that brain structural measures, such as cortical thickness and volumes, can vary along with the fluctuations of current symptoms(15).

However, identifying the neural associations of dynamic longitudinal features of depressive symptoms within a single well-powered imaging study has to date been challenging due to a lack of suitable data. In order to identify imaging correlates of depressive symptoms, a large imaging sample with repeated and consistently assessed measures of depressive symptoms is required. Such studies are resource intensive, take many years to complete and rarely provide the data required in sufficient numbers of participants. The UK Biobank Imaging Study (https://imaging.ukbiobank.ac.uk/) however is a rare exception and is by far the largest neuroimaging cohort with longitudinal depressive symptom data to date.

In the UK Biobank Imaging study, depressive symptoms were assessed on up to four separate occasions, across a time span of between 5.89 to 10.69 years. One depressive symptom assessment was conducted at the same occasion as the MRI imaging evaluation. Based on all available measurements, we generated four measures of depressive symptoms under two categories. The first category contained (i) a *single cross-sectional* assessment of depressive symptom severity at the same time as the imaging assessment, representing the current levels of symptoms of depression. The second category contains three measures derived from multiple assessments. These *longitudinal* measures include (ii) the longitudinal slope of depressive symptoms within an individual over time up until the imaging assessment, this was used as a proxy for assessing the longitudinal course of depressive symptoms over time, (iii) the mean level of depressive symptoms averaged over all measures and (iv) the standard deviation of depressive symptoms, as a measure of within-participant variability over time.

In the present study, we first investigated the associations between these measures of depressive symptoms and white matter microstructure, due to growing evidence of an association between mood disorders and reductions of white matter microstructure in the limbic system, especially in the thalamic radiations(16; 17). These networks contain important tracts involved in emotional processing(18) and regulation(19), deficits in which are typically associated with the onset and severity of MDD(20). In the present study, N = 19,345 people with diffusion tensor imaging data were included in order to test the association between white matter microstructure and the cross-sectional and longitudinal measures of depressive symptoms(21). Secondly, to explore potential differential contributions, we used step-wise regression models to test which brain regions in particular were associated with the cross-sectional measure of current symptoms, over and above those associated with longitudinal measures, and conversely, which are particularly sensitive to longitudinal measures, over and above those associated with current symptoms from the cross-sectional measures. Finally, since these dynamic features may reflect differential clinical and behavioural profiles, we also tested correlations between the above cross-sectional and longitudinal measures themselves, and their associations with 12 potentially MDD-relevant non-imaging behavioural, demographic and cognitive measures.

## Methods

### Participants

UK Biobank initially recruited 0.5M people across the United Kingdom(22), and has had an ongoing programme of follow-up studies, many of which have repeated the behavioural assessments conducted at baseline. In addition, UK Biobank has begun a brain imaging study of 100K individuals, as part of a wider battery of imaging investigations (21). For the current analyses, the most recent release of imaging data was used (October 2018). This sample included N=19,345 individuals who provided data that passed the quality check performed by UK Biobank imaging team after data pre-processing(23). In this sample, the mean age of participants was 63.06 years (SD = 7.44) and 47.14% were men. We then conducted further data quality control of the remaining participants, removing outliers(16; 17). Finally, the imaging data was merged with other relevant phenotypic data, including measures of depressive symptoms (steps listed below and presented in Figure S1).

UK Biobank data acquisition was approved by Research Ethics Committee (reference 11/NW/0382). The analysis and data acquisition for the present study were conducted under application #4844. Written consent was obtained for all participants. All the imaging preprocessing was undertaken according to the protocol provided by UK Biobank (https://biobank.ctsu.ox.ac.uk/crystal/docs/brain_mri.pdf).

### Depressive symptoms

Depressive symptoms were measured by a 4-item physical health questionnaire (PHQ-4)(24). PHQ-4 has an AUC (area under the curve) of 0.79 for its correlation with depression diagnosis(25). In other work, this measure was significantly associated with measures of disability(26) as well as risk factors for depression(24; 27). See more details in the URL: http://biobank.ctsu.ox.ac.uk/crystal/label.cgi?id=100060, and supplementary methods.

PHQ-4 was assessed repeatedly up to four times. Time points included: (a) the first assessment visit (2006-2010, N=19,231), (b) a repeat visit on a sub-sample (2012-2013, N=4,535), (c) the imaging visit (2014-2017, N =19,113) and finally the (d) online follow-up (2015-2018, N =14,155). Further details can be found on the UK Biobank website: http://biobank.ctsu.ox.ac.uk/crystal/label.cgi?id=100060.

Based on the repeated PHQ-4 measures, we generated four measures of depressive symptoms (Figure 1, S2): (i) a single PHQ-4 score measure acquired at the time of imaging assessment, representing the current depressive symptoms. Three other measures were generated based on multiple assessments, which included (ii) the estimated longitudinal slope of depressive symptoms from the initial recruitment up until the imaging assessment, using a linear growth curve model, positive values indicating a relative increased (i.e. worsening) progression over time, (iii) the mean and (iv) variability of depressive symptoms across all available assessments, where mean depression level was the average of PHQ-4 over two or more time points, and variability of depressive symptoms was the standard deviation of PHQ-4 scores over a minimum of three time points. Details of the growth curve model estimation are detailed in the supplementary methods. Descriptive statistics for the above PHQ-4 measures are presented in Figure S2 and Table S1.

**Figure 1.**
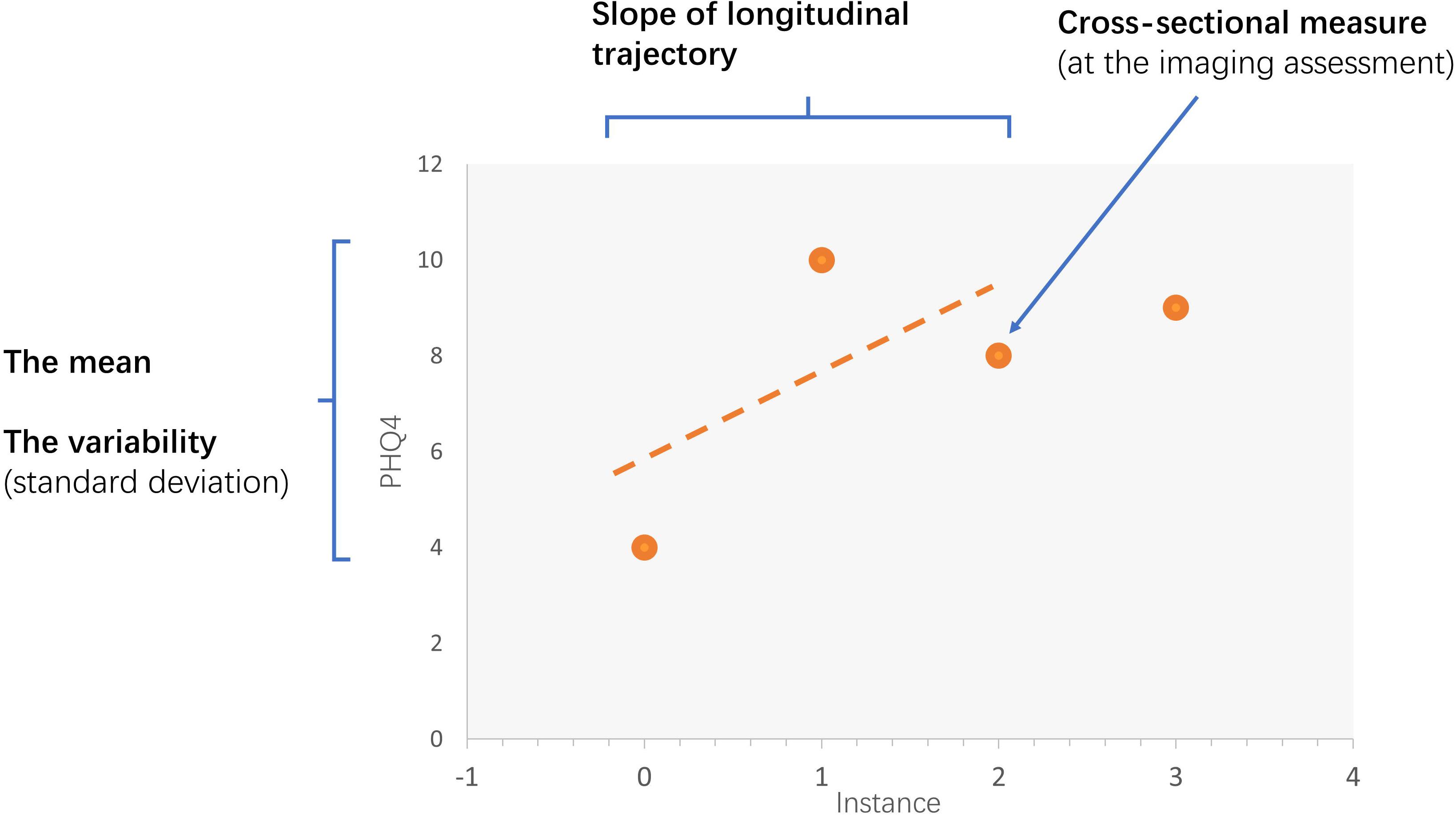
The measures for depressive symptoms generated for this study. There were four measures generated: (1) cross-sectional measurement for depressive symptoms acquired with imaging assessment, (2) linear growth curve denoting longitudinal trajectory of depressive symptoms derived from three time-points up until the imaging assessment, (3) mean of depressive symptoms generated based on at least two multiple assessments, and finally (4) variability of depressive symptoms which was the standard deviation of at least three time-points.

### Imaging data

We used the quality-controlled imaging-derived phenotypes (IDPs) from the diffusion tensor imaging (DTI) assessments released by the UK Biobank Imaging Study. Data acquisition, data pre-processing, estimation of white matter microstructure and quality check after the above steps were conducted by UK Biobank using a standard protocol described in the Primary Brain Imaging Documentation (URL: https://biobank.ctsu.ox.ac.uk/crystal/docs/brain_mri.pdf) and in two published protocol papers(21; 23). The procedures are described in brief below.

All imaging data was acquired using a 3T Siemens Skyra (software platform VD13) machine, using a standard (“monopolar”) Stejskal-Tanner pulse sequence. FSL packages were used for data pre-processing and microstructure estimation(28). Pre-processing included correction for eddy currents, head-motion, and gradient distortion, using the Eddy tool(29). Two DTI microstructure measures were estimated after pre-processing. Fractional anisotropy (FA) and mean diffusivity (MD) were generated using DTIFIT(30).

The processed data was then fed into AutoPtx package from FSL (https://fsl.fmrib.ox.ac.uk/fsl/fslwiki/AutoPtx), which uses probabilistic-tractography based methods to map 27 major tracts over the whole brain(21). The processed tracts included 12 bilateral and 3 unilateral tracts (Figure S1 and supplementary methods). FA data was used for mapping and the masks of tracts for each individual were used to locate the tracts on MD. Weighted means of DTI measures for each tract were generated.

Three newly developed neurite orientation dispersion and density imaging measures (known as the ‘NODDI’ measures) were also generated using AMICO tool as supplementary measures to the classic DTI measures(31). These measures depict additional sources of variation to FA and MD such as neurite density, extracellular water proportion, and morphology of tract organization. For brevity, these findings are not presented in the current manuscript, but for completeness, they are detailed in supplementary materials (Figures S4 and S5, Tables S4 and S6).

### MDD-related behavioural data

We also examined the association of each depressive symptom measure with an additional 12 potentially-relevant behavioural, demographic and cognitive measures that have been found to be associated with major depressive disorder or depressive symptoms in previous studies(32–34), The 12 measures included neuroticism, self-declared age of onset for depression, sociodemographic variables (household income, educational attainment and Townsend index), lifestyle measures (insomnia and smoking status), physical measures (body mass index and hand grip strength). Finally, we also examined the association of each symptomatic variable with measures of recent pain and general cognitive functions (a general ‘g’ factor generated by the first unrotated principal component of several cognitive tasks). More details are provided in the methods section within supplementary materials.

## Statistical methods

### Associations between measures of depressive symptoms and white matter microstructure

Before analysis was performed, outliers were removed (16). This was achieved by performing separate PCA for each DTI measure on the overall sample of N=19,345, those who were outside of +/- 3 standard deviations from mean were removed(16). This resulted in a maximum of N=19,262 participants remaining (see Figure S2) for further analysis.

We tested associations between each measure of depressive symptoms and the white matter microstructure measures with increasing level of regional detail: first using whole brain metrics, then for three tract subsets (using categories of: association/commissural fibres, thalamic radiations and projection fibres (see Figure S1)(16; 35)), and finally examining individual tracts separately. Indices for the whole brain and tract categories were derived as previous from the scores of the first un-rotated principal component for each microstructural metric. These are denoted gTotal, gAF, gTR and gPF (Figure S4 and Table S2) (16; 35).

The “glm” function in R was used to test for associations between each symptomatic measure and each unilateral tract. We used the R package “lme” for the analysis of bilateral tracts (where hemisphere was controlled for and each tract modelled as a repeated measure(36)). Age, age^2^, gender, head position in the scanner (on x, y and z axis), and MRI site were set as covariates. Other covariates included smoking status and alcohol consumption before the time of imaging assessment to control for depression-related behavioural patterns that may influence brain structure. We also adjusted for recent stressful life events within two years before the imaging assessment in order to have more accurate estimations of the associations related to inherent mood variability that may contribute variance to the measures of depressive symptoms. Each of the covariates are described in supplementary materials. For completeness, we also report results that did not control for smoking status, alcohol consumption, stressful life events in the supplementary material (Figure S6).

### Identifying the separate white-matter associations of trait and state measures of depressive symptoms

Cross-sectional measures of symptom severity, as an index of current ‘state’, and the longitudinal mean and variability, as indices of the ‘trait’, may be expected to be correlated, but also potentially distinctive. Therefore, we investigated which regions were more specifically associated with current ‘state’ by evaluating white matter associations with depressive symptom severity measured at the time of the imaging assessment, while adjusting for the longitudinal mean and variability symptom estimates. We also explored the reverse, testing which regions were associated with longitudinal ‘trait’ features (mean and variability) over and above the single cross-sectional measure of current symptoms obtained at the time of the imaging assessment.

To directly test this hypothesis of differential neurobiological associations with state and trait features, we utilised a step-wise regression method, using the ‘anova’ function in R. This tested whether the added independent variables significantly contributed to a reduction in the residual term of the model. First, we undertook a comparison of models to determine the unique contribution of cross-sectional symptom severity associations with imaging measures. Here we define:

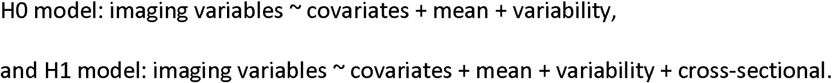

By comparing the H0 and H1 models, a significant reduction of residual in the H1 model compared with the H0 model would indicate an identifiable contribution of the cross-sectional symptoms measure to the model, and therefore a ‘state’-specific association between depressive symptoms and white matter microstructure.

Similarly, in order to test which imaging variables, the longitudinal measures contribute significantly over and above the cross-sectional measures, H0 and H1 models were defined as:

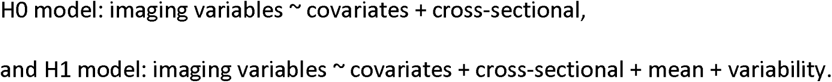

Results for the above analyses can be found in Figure 3 and Table S7-8.

**Figure 2.**
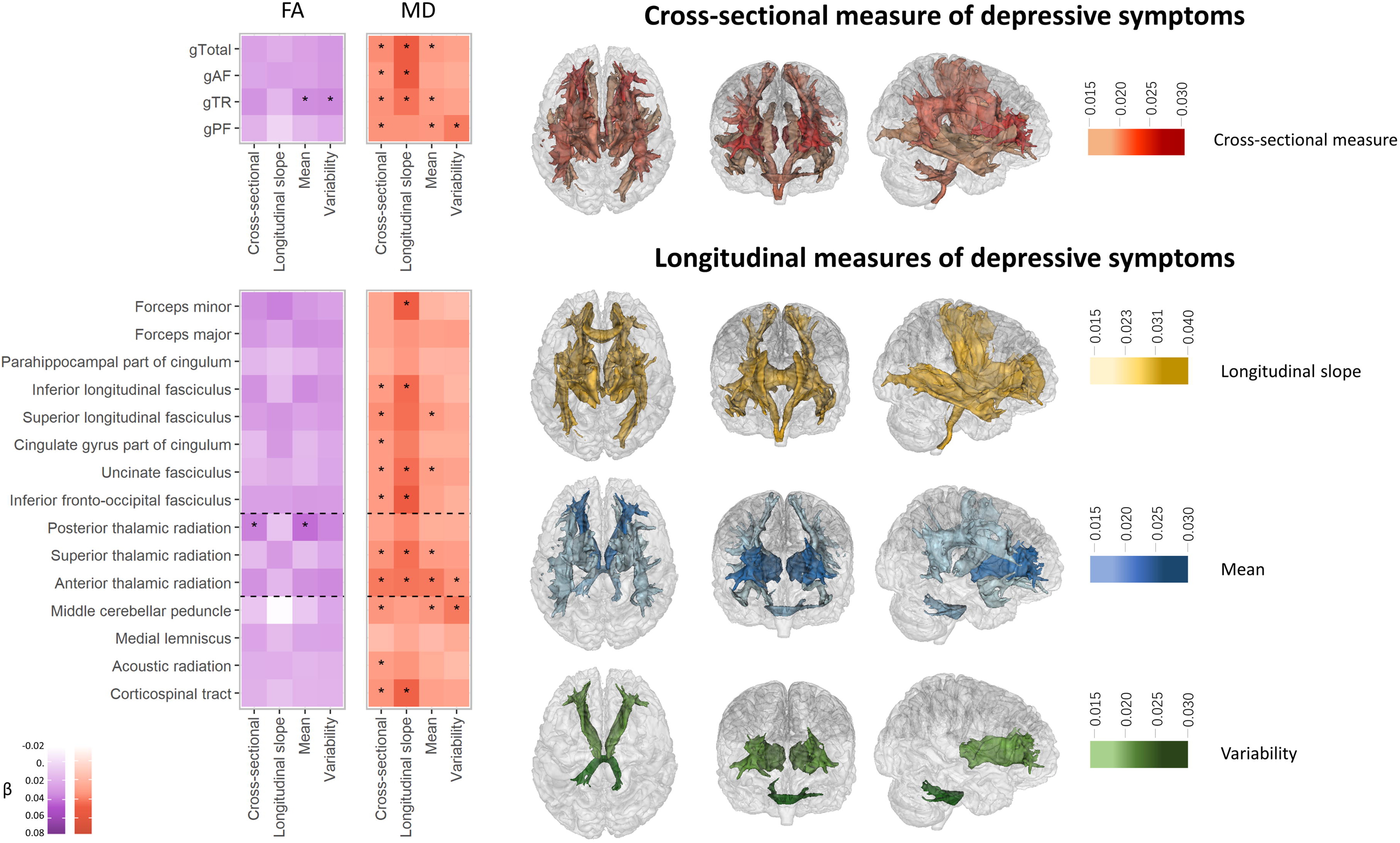
Associations between cross-sectional depressive symptoms and dMRI (heatmap) and the map for significant regions (brain map). FA=fractional anisotropy, MD=mean diffusivity. For the heatmap, colour depth represents the standard effect size of a measure. As FA has negative direction with MD, here in this figure, the effect sizes for FA was reversed (×-1). The results were separated in two sections. The upper sections were the results for g measures and the lower sections showed the results of individual tracts. To aid comprehension, the lower part where results of tracts were shown, checks were divided into three categories by dashed lines as the tracts were in different subsets, i.e. association fibres, thalamic radiations and projection fibres (see Methods). Significant associations after FDR correction (p_corr_<0.05) were marked with an asterisk. For the brain map, significant tracts were shown in red for the ones associated with cross-sectional measure, yellow for the ones with longitudinal slope, and orange for both. Significant tracts were shown in blue if associated with the mean, light green for variability, and turquoise for both.

**Figure 3.**
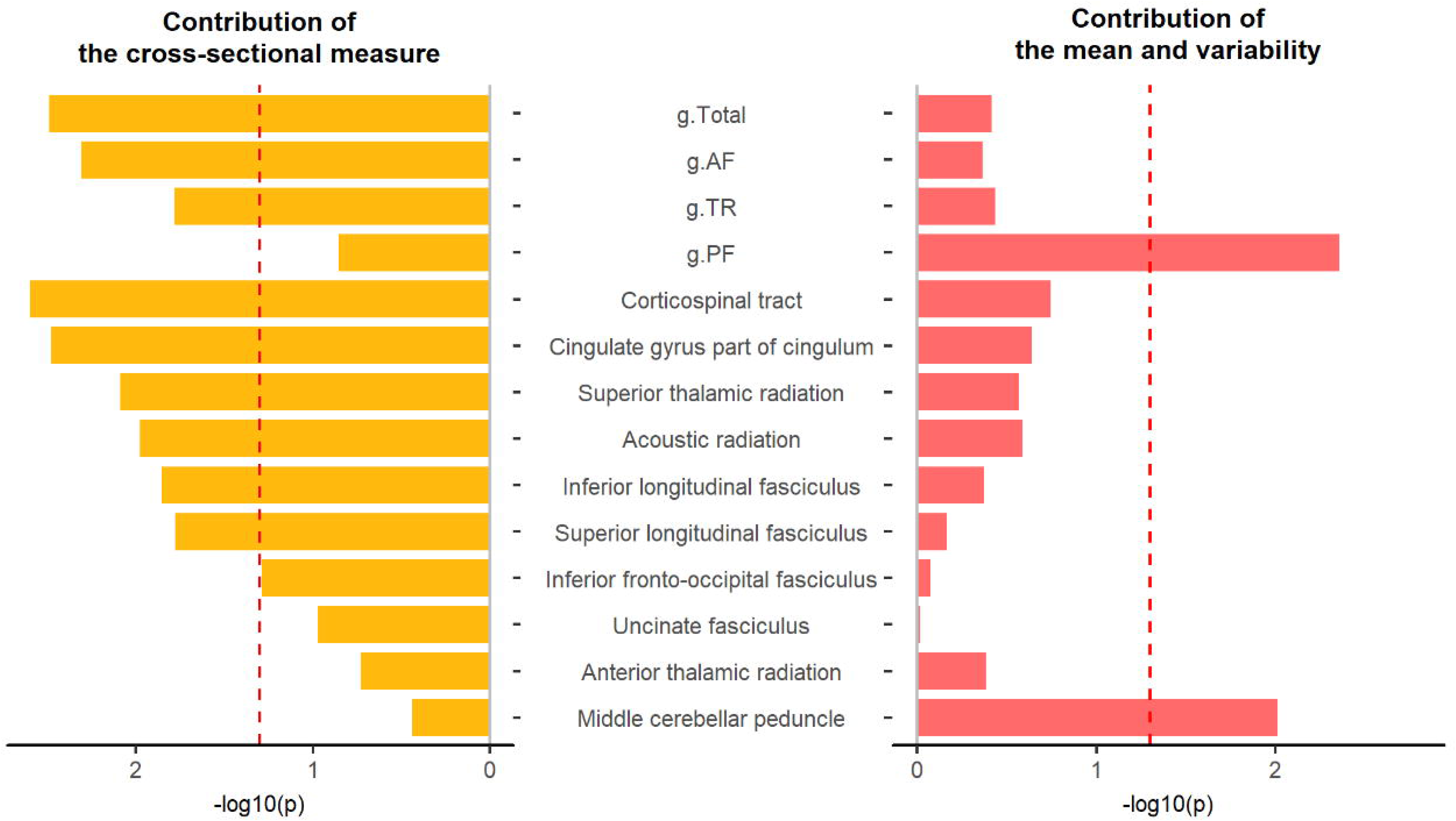
Contributions by the cross-sectional measure and by the mean and variability of depressive symptoms on MD (mean diffusivity). The y axis represents the variables that were found associated with cross-sectional depressive symptoms. The x axis represents the -log10(p) for whether adding relevant measure(s) of depressive symptoms adds significantly more variance explained. For the left panel, an H1 model with independent variables that include the mean, variability and cross-sectional measure was compared with an H0 model with only the mean and variability were the dependent variables. Bars to the left of the red dashed line (the p=0.05 line) indicate a significant contribution of the cross-sectional measure. Vice versa for the right panel, in which the bars to the right the red dashed line indicate a significant contribution of the mean and variability for MD in these neuroimaging variables. For measures of white matter microstructure, g.Total = global variance of MD, g.AF = g measure for association/commissural fibres, g.TR = g measure for thalamic radiations, g.PF = g measure for projection fibres.

### Associations between measures of depressive symptoms and behavioural variables

We first analysed the correlations within the four measures of depressive symptoms themselves, and then conducted analysis on the associations between these measures with other behavioural variables. For the associations between measures of depressive symptoms and behavioural variables, they were tested controlling for age, age^2^, sex and MRI site using glm function in R version 3.2.3 (see supplementary material). P values were false discovery rate (FDR)-corrected using “p.adjust” function in R (q-value<0.05) applied for 4 (symptom measures) *12 (behavioural variables) =48 tests. All effect sizes throughout the manuscript are standardised. For the logistic regressions conducted on binary dependent variables, the effect sizes were reported as log-transformed odds ratio.

## Results

### Associations between measures of depressive symptoms and white matter microstructure

The anterior thalamic radiation was the only structure that was significantly associated with all four measures of depressive symptoms (β ranged 0.028 to 0.030, p_corr_<0.049), See Figure S2 and Table S3-4. The largest effect sizes for the white matter associations were however shown with measures longitudinal trajectory (β ranged 0.030 to 0.040 for significant associations, p_corr_<0.049). Cross-sectional, mean and variability of depressive symptoms generally had lower effect sizes than the longitudinal slope associations (where βs ranged from 0.015 to 0.029 for significant associations, p_corr_<0.044). The main results for each of the four measures of depressive symptoms are described in detail below. Results without controlling for smoking status, alcohol consumption and stressful life events demonstrated similar patterns of results to the main findings controlling for these factors (see supplementary materials).

### Cross-sectional measure of depressive symptoms

Higher general MD (g) in all tract categories including the general variance over all tracts (gTotal), and tracts separated into the categories of association/commissural fibres (gAF), thalamic radiation (gTR) and projection fibres (gPF) were positively associated with cross-sectional depressive symptoms (β ranged from 0.019 to 0.024, p_corr_ ranged from 0.009 to 0.002).

MD in a total of ten individual tracts were found to be associated with cross-sectional measures of depressive symptoms. Higher cross-sectional symptom severity was positively associated with higher MD in the acoustic radiation (β=0.015, p_corr_=0.041), anterior thalamic radiation (β=0.028, p_corr_=1.47×10^−4^), inferior longitudinal fasciculus (β=0.016, p_corr_=0.040), inferior fronto-occipital fasciculus (β=0.017, p_corr_=0.038), uncinate fasciculus (β=0.016, p_corr_=0.038), superior thalamic radiation (β =0.020, p_corr_=0.011), corticospinal tract (β =0.020, p_corr_=0.030), superior longitudinal fasciculus (β =0.022, p_corr_=0.013), cingulate gyrus part of cingulum (β =0.017, p_corr_=0.038), and middle cerebellar peduncle (β =0.020, p_corr_=0.038).

For FA, no significant associations were found for the global measures (whole brain and tract categories) to be associated with the cross-sectional symptom severity (p_corr_>0.051). For the individual tracts, only FA in posterior thalamic radiation was associated with higher cross-sectional measure (β=-0.024, p_corr_=0.011).

### Longitudinal measures of depressive symptoms: longitudinal slope, the mean and variability

Greater increase of longitudinal symptomology over time was positively associated with gTotal, gAF and gTR for MD (β ranged from 0.031 to 0.040, p_corr_ ranged from 0.025 to 0.009).

Both higher mean and greater variability of depressive symptoms were found to be positively associated with higher MD in gPF (mean: β =0.020, p_corr_=0.009, variability: β=0.029, p_corr_=0.002). Additional associations with the mean level of depressive symptoms were seen in global MD (β =0.018, p_corr_=0.013) and MD in the subset of gTR (β =0.018, p_corr_=0.009).

Tract-specific analysis showed that greater longitudinal increase of depressive symptoms was positively associated with higher MD in the anterior thalamic radiation (β=0.030, p_corr_=0.049), corticospinal tract (β=0.035, p_corr_=0.038), inferior fronto-occipital fasciculus (β=0.038, p_corr_=0.030), inferior longitudinal fasciculus (β=0.033, p_corr_=0.040), superior thalamic radiation (β=0.031, p_corr_=0.040), uncinate fasciculus (β=0.032, p_corr_=0.038) and forceps minor (β=0.037, p_corr_=0.038).

For mean and variability measures, MD in individual tracts that were positively associated with both higher mean and variability of depressive symptoms were seen in the anterior thalamic radiation (mean: β=0.029, p_corr_=1.47×10^−4^, variability: β=0.020, p_corr_=0.030), and middle cerebellar peduncle (mean: β=0.019, p_corr_=0.038, variability: β=0.029, p_corr_=0.008). Additional associations that were found for the mean of depressive symptoms were shown for higher MD in superior longitudinal fasciculus (β=0.017, p_corr_=0.042), uncinate fasciculus (β=0.015, p_corr_=0.044) and superior thalamic radiation (β=0.016, p_corr_=0.038).

FA in gTR was negatively associated with mean level and within-participant variability of depressive symptoms over time (β=-0.021 and -0.022, p_corr_=0.045 and 0.045 respectively). For individual tracts, the only association was found between FA in posterior thalamic radiation and mean depressive symptoms (β=-0.029, p_corr_=7.85×10^−4^). No significant association between tract-specific FA variation and longitudinal slope or variability of depressive symptoms were found (p_corr_ >0.055).

### Assessing the associations of current symptoms after adjustment for longitudinal measures, and vice versa

As shown in Figure 3, a significant additional contribution of the cross-sectional symptom measure, over and above the longitudinal measures, was found for MD in gTotal, gAF and gTR (F statistics for H0-H1 model comparison ranged from 5.75 to 8.66, p ranged from 0.017 to 0.003). Additional contribution by the cross-sectional symptom measure was also shown for individual tracts including the superior longitudinal fasciculus, superior thalamic radiation, inferior longitudinal fasciculus, corticospinal tract, acoustic radiation and cingulate gyrus part of cingulum (F ranged from 5.73 to 9.12, df = 1, p ranged from 0.017 to 0.003).

Conversely, MD in gPF showed a significant additional contribution from the mean and variability measures over the cross-sectional measure (F= 5.43, df=2, p = 0.004). For individual tracts, significant additional variance contributed by the mean and variability of depressive symptoms was found in middle cerebellar peduncle (F = 4.63, df=2, p = 0.010).

### Associations between measures of depressive symptoms and behavioural traits

The cross-sectional measure of depressive symptoms was positively and significantly correlated with all three longitudinal measures of depressive symptoms with r = 0.457, 0.840 and 0.478 (all p < 10^−16^) for the correlations with longitudinal slope, mean and variability of depressive symptoms respectively. A correlation matrix between the measures can be found in Table S1.

For the depression-related behavioural variables, contrary to the imaging results, the longitudinal trajectory of symptoms over time had the lowest measures of associations (absolute β ranged from 0.026 to 0.081, p_corr_<0.043 for significant associations) compared all other three measures of depressive symptoms. The largest effects were for overall mean symptom severity (absolute β ranged from 0.048 to 0.216, p_corr_<3.81×10^−6^). There was no clear pattern of difference between associations with current symptoms versus the longitudinal mean and variability for these behavioural variables. There were however indications of a similar pattern of the strongest associations for all of these three measures of depression severity with neuroticism, insomnia and pain (cross-sectional: absolute β ranged from 0.195 to 0.524, p_corr_<1×10^−30^ ; the mean: absolute β ranged from 0.216 to 0.567, p_corr_<1×10^−30^; and variability: absolute β ranged from 0.145to 0.399, p_corr_<1×10-30). Other significant associations are detailed in Table 1.

**Table 1.**
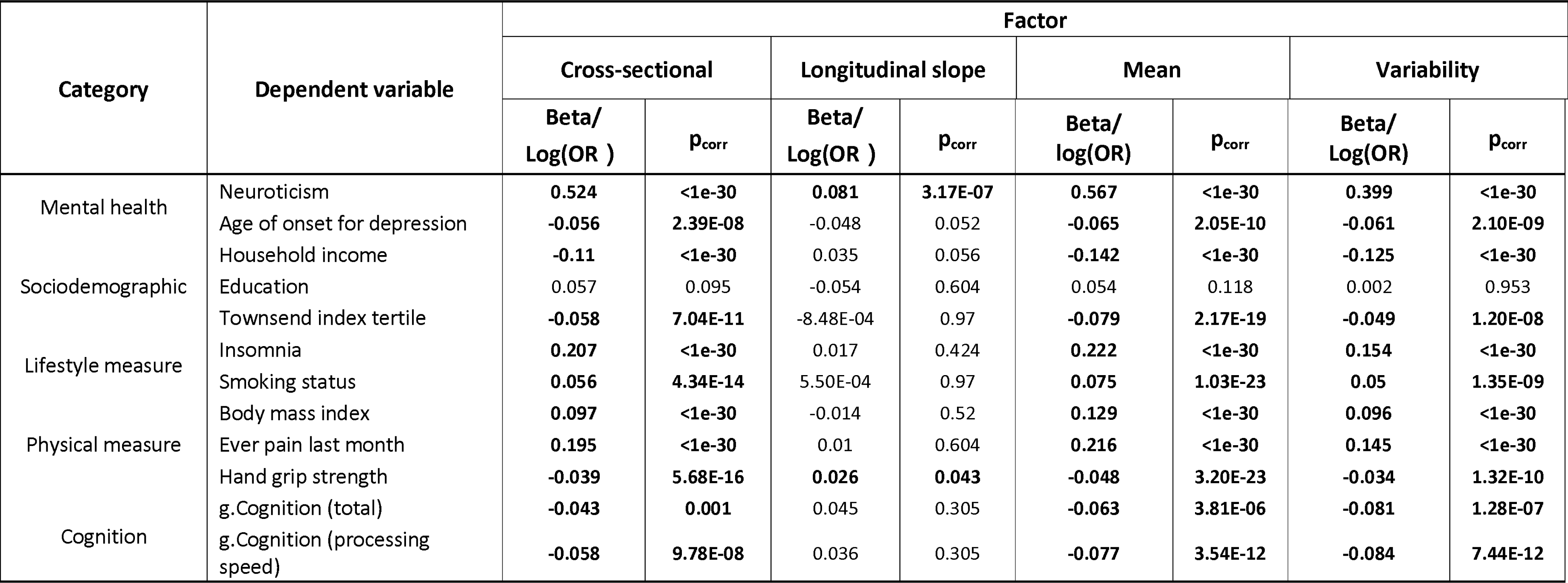
Associations between measures of depressive symptoms and behavioural variables. The greatest effect sizes across all four measures were highlighted in red for each line. Log (OR) = log (odds ratio).

## Discussion

In the current investigation we report novel patterns of association between four measures of cross-sectional and longitudinal depressive symptom severity with decreased WM microstructural integrity. Decreased white matter microstructure in the anterior thalamic radiation demonstrated significant associations across all four measures of depressive symptoms (for MD; β ranged 0.028 to 0.030). The strongest white matter associations were found for variables relating to increasing longitudinal symptom severity. Measures of current symptom severity (cross-sectional measures) were particularly associated with decreased white matter integrity in association fibres and thalamic radiations (for MD; β=0.015 to 0.039). Associations with higher mean and variability of depressive symptoms over time however showed associations primarily in projection fibres (for MD; β=0.019 to 0.029). Contrary to the imaging findings, the non-imaging variables, in particular neuroticism, insomnia and pain, were associated with mean and variability of symptoms over time, as well as with current symptoms, rather than for longitudinal change in symptoms.

Stable and transient conditions, referred to as ‘trait’ and ‘state’ in other research contexts, are two related and yet distinctive features contributing to individual differences in mood conditions. Recent population-based genetic and predictive modelling studies have revealed that stable manifestations of emotional problems typically have a higher heritability than more transient features(37). Also longitudinal measures, including variability, had a higher predictive power relating to severe forms of behaviour like suicide attempts(38). Hence, understanding biological basis of these potentially distinctive features would have important implications(39; 40).

Deficits in anterior thalamic radiation in particular showed associations with all four measures of depressive symptoms. The thalamic radiation tract subset was also consistently associated with all measures of depressive symptoms (nominally significant for variability measure and significant for all other measures). This shared mechanism between measures indicates the importance of thalamic system, particularly fronto-thalamic connectivity, in depressive symptomatology. Our results could therefore explain why thalamic radiations are one of the most replicated findings in either life-time or cross-sectional case-control studies of MDD (16; 41; 42). There are two potential reasons for the above results. First, deficits in microstructure of the thalamic radiation may be particularly susceptible to the influence by early life factors (43; 44), and second, they may relate to genetic predisposition to depression (17). Although the observational data in the present study did not allow for causal inferences, in future studies, thalamic radiations may be a possibly strong candidate as a causal biomarker for illness.

Associations between cross-sectional measure with fibres in the subset of association fibres were particularly significant after correcting for longitudinal measures, which indicate that these regions are particularly sensitive to temporary variations. Deficits of microstructure in association fibres were also associated with worsening depressive symptoms over time. Association fibres and connections to the prefrontal cortex have been repeatedly found to be associated executive cognition(35; 45; 46) and closely related to psychological resilience(47). Abnormalities in these tracts could therefore contribute to both temporary depressive status and longitudinal decline in mental well-being(48).

Projection fibres were particularly associated with mean and variability of depressive symptoms. Microstructure in this white matter subset is well-known for being related to motor response and processing speed. Therefore, the results suggest that higher mean and variability of depressive symptoms may be related to cognitive decline and decline of psychomotor abilities(49; 50).

We note importantly that most of the current results were found for MD rather than FA. Despite of the differences in level of significance, MD and FA presented similar directions of effects and the scales of effect were similar for the most robust findings, especially in thalamic radiations (Table S3). The discordance between MD and FA may be rooted in the differential sensitivity of MD and FA to a variety of complex degenerative processes. Changes in FA, which for example could result from increased transverse diffusion due to myelin and axonal disruption(51), may be masked by co-occurring processes such as fibre reorganisation and glial reactivity. In such instances, where all three eigenvectors of the diffusion tensor experience proportional change, it is plausible that MD would offer greater sensitivity(52). Notably, MD also reportedly exhibits greater sensitivity than FA to other traits, such as ageing(35).

In the present study, we used a very large imaging sample of over 18,900 people. Though the sample size of longitudinal change of depressive symptoms over four time points was much lower (∼4,000), it is still much larger than most neuroimaging studies, especially considering the data is longitudinal and covers up to ten years. All of this provides high statistical power to reliably detect modest associations(53). However, a limitation for the present study is that the time lag between adjacent assessments may vary per participant from three years to six years. For this reason, we also adjusted for this difference, by controlling for the age of each time point, and tested for the fit of the growth curve models (see supplementary methods).

Our results provide evidence that deficits in WM microstructure are related to greater longitudinal deterioration, current symptom severity, mean and variability of depressive symptoms, with different as well as overlapping regional patterns. Further mechanistic insights underlying the relationship between changes in neurobiology and changing symptoms will dependent on availability of future large-scale longitudinal neuroimaging datasets, along with the availability of methods and tools able to test for causal inferences.

## Acknowledgments

This study is supported by a Wellcome Trust Strategic Award “Stratifying Resilience and Depression Longitudinally” (STRADL) (Reference 104036/Z/14/Z).

This research was conducted using the UK Biobank Resource under approved project #4844. We thank the UK Biobank participants for their participation, and the UK Biobank team for their work in collecting and providing these data for analyses. Part of the work was undertaken in The University of Edinburgh Centre for Cognitive Ageing and Cognitive Epidemiology (CCACE), funding from the Biotechnology and Biological Sciences Research Council (BBSRC) and Medical Research Council (MRC) is gratefully acknowledged.

XS receives support from China Scholarship Council (201506040037). HCW is supported by a JMAS SIM fellowship from the Royal College of Physicians of Edinburgh and by an ESAT College Fellowship from the University of Edinburgh. Author SRC was supported by the Medical Research Council grants MR/M013111/1 and MR/R024065/1, by the Age UK-funded Disconnected Mind project, and a National Institutes of Health research grant (R01AG054628). TER is supported by funding from the Wellcome Trust 4-year PhD in Translational Neuroscience [108890/Z/15/Z]. AMM is supported by MRC Mental Health Pathfinder Grant MC_PC_17209.

## References

1. Vos T, Flaxman AD, Naghavi M, Lozano R, Michaud C, Ezzati M, et al. (2012): Years lived with disability (YLDs) for 1160 sequelae of 289 diseases and injuries 1990-2010: A systematic analysis for the Global Burden of Disease Study 2010. Lancet. 380: 2163–2196.

2. Sullivan PF, Neale MC, Kendler KS (2000): Genetic Epidemiology of Major Depression: Review and Meta-Analysis. Am J Psychiatry. 157: 1552–1562.

3. Kessler RC, Berglund P, Demler O, Jin R, Koretz D, Merikangas KR, et al. (2003): The Epidemiology of Major. Am Med Assoc. 289: 3095–3105.

4. Whalley H, Harris M, Shen X, Gibson J, Lawrie S, McIntoshl A (2018): S130. Dissecting the Neuroimaging Phenotype of Major Depressive Disorder Based on Genetic Loading for Schizophrenia. Biol Psychiatry. 83: S398.

5. Howard DM, Clarke T-K, Adams MJ, Hafferty JD, Wigmore EM, Zeng Y, et al. (2017): The stratification of major depressive disorder into genetic subgroups David. bioRxiv. doi: 10.1130/G38594.1.

6. Liao Y, Huang X, Wu Q, Yang C, Kuang W, Du M, et al. (2013): Is depression a disconnection syndrome? Metaanalysis of diffusion tensor imaging studies in patients with MDD. J Psychiatry Neurosci. 38: 49–56.

7. Fava M, Kendler KS (2000): Major Depressive Disorder Review. Neuron. 28: 335–341.

8. Kendler KS, Gardner CO (2011): A longitudinal etiologic model for symptoms of anxiety and depression in women. Psychol Med. 41: 2035–2045.

9. van Eeden WA, van Hemert AM, Carlier IVE, Penninx BW, Giltay EJ (2019): Severity, course trajectory, and within-person variability of individual symptoms in patients with major depressive disorder. Acta Psychiatr Scand. 139: 194–205.

10. Cramer AOJ, Van Borkulo CD, Giltay EJ, Van Der Maas HLJ, Kendler KS, Scheffer M, Borsboom D (2016): Major depression as a complex dynamic system. PLoS One. 11: 1–20.

11. Malhi GS, Mann JJ (2018): Depression. Lancet. 392: 2299–2312.

12. Linden DEJ (2012): The Challenges and Promise of Neuroimaging in Psychiatry. Neuron. 73: 8–22.

13. Howard DM, Adams MJ, Shirali M, Clarke T-K, Marioni RE, Davies G, et al. (2017): Genome-wide association study of depression phenotypes in UK Biobank (n = 322,580) identifies the enrichment of variants in excitatory synaptic pathways. bioRxiv. 6268: 168732.

14. Wray NR, Ripke S, Mattheisen M, Trzaskowski M, Byrne EM, Abdellaoui, A., & Bacanu SA (2018): Genome-wide association analyses identify 44 risk variants and refine the genetic architecture of major depression. 50. doi:10.1038/s41588-018-0090-3.

15. Phillips JL, Batten LA, Tremblay P, Aldosary F, Blier P (2015): A Prospective, Longitudinal Study of the Effect of Remission on Cortical Thickness and Hippocampal Volume in Patients with Treatment-Resistant Depression. Int J Neuropsychopharmacol 1–9.

16. Shen X, Reus LM, Cox SR, Adams MJ, Liewald DC, Bastin ME, et al. (2017): Subcortical volume and white matter integrity abnormalities in major depressive disorder: Findings from UK Biobank imaging data. Sci Rep. 7: 1–10.

17. Barbu MC, Zeng Y, Shen X, Cox SR, Clarke T, Adams MJ, et al. (2018): Association of whole-genome and NETRIN1 signaling pathway-derived polygenic risk scores for Major Depressive Disorder and thalamic radiation white matter microstructure in UK Biobank. Biol Psychiatry Cogn Neurosci Neuroimaging. 44: 1–24.

18. Hall J, Whalley HC, McKirdy JW, Romaniuk L, McGonigle D, McIntosh AM, et al. (2008): Overactivation of Fear Systems to Neutral Faces in Schizophrenia. Biol Psychiatry. 64: 70–73.

19. Phillips ML, Ladouceur CD, Drevets WC (2008): A neural model of voluntary and automatic emotion regulation: Implications for understanding the pathophysiology and neurodevelopment of bipolar disorder. Mol Psychiatry. 13: 833–857.

20. Leppa JM (2006): Emotional information processing in mood disorderslll : a review of behavioral and neuroimaging findings. Curr Opin Psychiatry. 19: 34–39.

21. Miller KL, Alfaro-Almagro F, Bangerter NK, Thomas DL, Yacoub E, Xu J, et al. (2016): Multimodal population brain imaging in the UK Biobank prospective epidemiological study. Nat Neurosci. doi:10.1038/nn.4393.

22. Sudlow C, Gallacher J, Allen N, Beral V, Burton P, Danesh J, et al. (2015): UK Biobank: An Open Access Resource for Identifying the Causes of a Wide Range of Complex Diseases of Middle and Old Age. PLoS Med. 12: 1–10.

23. Alfaro-Almagro F, Jenkinson M, Bangerter NK, Andersson JLR, Griffanti L, Douaud G, et al. (2018): Image processing and Quality Control for the first 10,000 brain imaging datasets from UK Biobank. Neuroimage. 166: 400–424.

24. Batty GD, McIntosh AM, Russ TC, Deary IJ, Gale CR (2016): Psychological distress, neuroticism, and cause-specific mortality: Early prospective evidence from UK Biobank. J Epidemiol Community Health. 70: 1136–1139.

25. Khubchandani J, Brey R, Kotecki J, Kleinfelder JA, Anderson J (2016): The Psychometric Properties of PHQ-4 Depression and Anxiety Screening Scale Among College Students. Arch Psychiatr Nurs. 30: 457–462.

26. Kroenke K, Spitzer RL, Janet BW, Williams DSW, Lö B (2009): An Ultra-Brief Screening Scale for Anxiety and Depression: The PHQ– 4. Psychosomatics. 50: 613–621.

27. Löwe B, Wahl I, Rose M, Spitzer C, Glaesmer H, Wingenfeld K, et al. (2010): A 4-item measure of depression and anxiety: Validation and standardization of the Patient Health Questionnaire-4 (PHQ-4) in the general population. J Affect Disord. 122: 86–95.

28. Andersson JLR, Jenkinson M, Smith SM (2007): Non-linear optimisation. FMRIB technical report TR07JA1. In Pract.

29. Andersson JLR, Sotiropoulos SN (2015): An integrated approach to correction for off-resonance effects and subject movement in diffusion MR imaging. Neuroimage. 125: 1063–1078.

30. Anthofer JM, Steib K, Fellner C, Lange M, Brawanski A, Schlaier J (2015): DTI-based deterministic fibre tracking of the medial forebrain bundle. Acta Neurochir (Wien). 157: 469–477.

31. Daducci A, Canales-Rodríguez EJ, Zhang H, Dyrby TB, Alexander DC, Thiran JP (2015): Accelerated Microstructure Imaging via Convex Optimization (AMICO) from diffusion MRI data. Neuroimage. 105: 32–44.

32. Hagenaars SP, Harris SE, Davies G, Hill WD, Liewald DC, Ritchie SJ, et al. (2015): Shared genetic aetiology between cognitive functions and physical and mental health in UK Biobank (N = 112151) and 24 GWAS consortia. bioRxiv. 031120.

33. Clarke T-K, Hall LS, Fernandez-Pujals AM, MacIntyre DJ, Thomson P, Hayward C, et al. (2015): Major depressive disorder and current psychological distress moderate the effect of polygenic risk for obesity on body mass index. Transl Psychiatry. 5: e592.

34. Clarke TK, Lupton MK, Fernandez-Pujals AM, Starr J, Davies G, Cox S, et al. (2015): Common polygenic risk for autism spectrum disorder (ASD) is associated with cognitive ability in the general population. Mol Psychiatry. doi:10.1038/mp.2015.12.

35. Cox SR, Ritchie SJ, Tucker-Drob EM, Liewald DC, Hagenaars SP, Davies G, et al. (2016): Ageing and brain white matter structure in 3,513 UK Biobank participants. Nat Commun. 7: 1–34.

36. Chatfield C, Zidek J, Lindsey J (2010): An introduction to generalized linear models. Chapman and Hall/CRC.

37. Cheesman R, Depressive M, Working D, Consortium G (2018): Extracting stability increases the SNP heritability of emotional problems in young people. Transl Psychiatry. doi:10.1038/s41398-018-0269-5.

38. Melhem NM, Porta G, Oquendo MA, Zelazny J, Keilp JG, Iyengar S, et al. (2019): Severity and Variability of Depression Symptoms Predicting Suicide Attempt in High-Risk Individuals. JAMA Psychiatry. 15213: 1–11.

39. Huang H, Fan X, Williamson DE, Rao U (2011): White matter changes in healthy adolescents at familial risk for unipolar depression: a diffusion tensor imaging study. Neuropsychopharmacology. 36: 684–91.

40. Whalley HC, Papmeyer M, Sprooten E, Romaniuk L, Blackwood DH, Glahn DC, et al. (2012): The influence of polygenic risk for bipolar disorder on neural activation assessed using fMRI. Transl Psychiatry. 2: e130.

41. Coenen VA, Panksepp J, Hurwitz TA, Urbach H, Mädler B (2012): Human medial forebrain bundle (MFB) and anterior thalamic radiation (ATR): imaging of two major subcortical pathways and the dynamic balance of opposite affects in understanding depression. J Neuropsychiatry Clin Neurosci. 24: 223–36.

42. Frodl T, Carballedo A, Fagan AJ, Lisiecka D, Ferguson Y, Meaney JF (2012): Effects of early-life adversity on white matter diffusivity changes in patients at risk for major depression. J Psychiatry Neurosci. 37: 37–45.

43. Lai CH, Wu YT (2014): Alterations in white matter micro-integrity of the superior longitudinal fasciculus and anterior thalamic radiation of young adult patients with depression. Psychol Med. 44: 2825–2832.

44. Asato MR, Terwilliger R, Woo J, Luna B (2010): White matter development in adolescence: A DTI study. Cereb Cortex. 20: 2122–2131.

45. Voineskos AN, Rajji TK, Lobaugh NJ, Miranda D, Shenton ME, Kennedy JL, et al. (2012): Age-related decline in white matter tract integrity and cognitive performance: A DTI tractography and structural equation modeling study. Neurobiol Aging. 33: 21–34.

46. Zheng Z, Shemmassian S, Wijekoon C, Kim W, Bookheimer SY, Pouratian N (2014): DTI correlates of distinct cognitive impairments in Parkinson’s disease. Hum Brain Mapp. 35: 1325–1333.

47. Walsh ND, Williams SCR, Brammer MJ, Bullmore ET, Kim J, Suckling J, et al. (2007): A Longitudinal Functional Magnetic Resonance Imaging Study of Verbal Working Memory in Depression After Antidepressant Therapy. Biol Psychiatry. 62: 1236–1243.

48. Bachmann RF, Schloesser RJ, Gould TD, Manji HK (2005): Mood stabilizers target cellular plasticity and resilience cascades: Implications for the development of novel therapeutics. Mol Neurobiol. 32: 173–202.

49. Bammer R, Turken U, Whitfield-Gabrieli S, Gabrieli JDE, Dronkers NF, Baldo J V. (2008): Cognitive processing speed and the structure of white matter pathways: Convergent evidence from normal variation and lesion studies. Neuroimage. 42: 1032–1044.

50. Behrens TEJ, Menke RA, Gass A, Matthews PM, Kindlmann G, Whitcher B, et al. (2010): DTI measures in crossing-fibre areas: Increased diffusion anisotropy reveals early white matter alteration in MCI and mild Alzheimer’s disease. Neuroimage. 55: 880–890.

51. Jones DK, Knösche TR, Turner R (2013): White matter integrity, fiber count, and other fallacies: The do’s and don’ts of diffusion MRI. Neuroimage. 73: 239–254.

52. Acosta-Cabronero J, Williams GB, Pengas G, Nestor PJ (2010): Absolute diffusivities define the landscape of white matter degeneration in Alzheimer’s disease. Brain. 133: 529–539.

53. Smith SM, Nichols TE (2018): Statistical Challenges in “Big Data” Human Neuroimaging NeuroView. Neuron. 97: 263–268.

