## Supplementary materials for "White matter microstructure and its relation to longitudinal measures of depressive symptoms in mid-late life"

Shen *et al.*

**Table of contents**

Supplementary methods

Supplementary results

Figure S1-7.

Table S1-6.

**Supplementary methods**

**PHQ-4 questionnaire**

PHQ4 questions include: “Frequency of depressed mood in last 2 weeks”, “Frequency of unenthusiasm/disinterest in last 2 weeks”, “Frequency of tenseness/restlessness in last 2 weeks” and “Frequency of tiredness/lethargy in last 2 weeks”. This questionnaire assesses depression-related symptoms within a 2-week timeframe. The sum of the score was calculated to indicate depressive symptoms.

The mean time lag between the first and second occasion was 4.29 years with a standard deviation of 0.94 years. Between the second and third occasion, mean time lag was 3.32 years with a standard deviation of 1.19 years. Between the third and the final occasion, mean time lag was 0.003 years with a standard deviation of 1.10 years.

For each measure of depressive symptoms derived from cross-sectional assessments, the mean time lag of mean level of depressive symptoms was 8.04 years (sd=1.22 years), the variability of depressive symptoms was 8.09 years (sd=1.14 years), and slope of longitudinal trajectory was 8.26 years (sd=1.11 years). The correlations between longitudinal measures and time lag was very small and therefore this variable was not included in the main model (r ranged from 2.41×10^-4^ to 0.027).

**dMRI measures**

The processed tracts included 12 bilateral tracts that has a value for each brain hemisphere (acoustic radiation, anterior thalamic radiation, cingulate gyrus part of cingulum, corticospinal tract, inferior fronto-occipital fasciculus, inferior longitudinal fasciculus, medial lemniscus, parahippocampal part of cingulum, posterior thalamic radiation, superior longitudinal fasciculus, superior thalamic radiation and uncinate fasciculus) and 3 unilateral tracts (forceps major, forceps minor and middle cerebellar peduncle).

**Depressive symptoms**

For the growth curve model, we used the ‘growth’ function from the ‘lavaan’ package (<http://lavaan.ugent.be/tutorial/growth.html>) in R(1). Scaled age at each assessment was controlled for. The growth curve model showed good fit to the data (CFI = 0.988, TLI = 0.984, RMSEA = 0.031, SRMR = 0.014, Chi-square (17) = 103.563 with a p<0.001). Longitudinal change within the whole population was in a negative direction but did not reach to significance (β = -0.094, p = 0.164). Both the intercept (β= 7.366, p < 0.001) and variance (β=1.521, p < 0.001) of the mean slope of growth curve model were significant. Each individual’s slope of longitudinal trajectory was estimated for further analysis by using the ‘predict’ function in ‘lavaan’.

**Covariates**

In addition to age, age^2^ and gender, we also included scanner positions for all three axis, alcohol consumption, smoking status and stressful life events. The covariates except for age and gender will be explained in detail below. All the covariates described here were acquired with the imaging assessments. See also Table S1, S5 and S6.

The scanner position was used for controlling for systematic change in the static magnetic field (the last four fields in <http://biobank.ctsu.ox.ac.uk/crystal/label.cgi?id=110>). These proxies for scanner position showed minimal correlation with our white matter phenotypes, but in order to achieve a better estimated model, we chose to include them in our models.

Alcohol consumption was self-reported weekly consumption which was used in a published paper on the overall UK Biobank sample of about 500k people (2). We used a slightly different approach to exclude impossible numbers. In the referenced study, they excluded values over 5 standard deviations from mean, and we employed their values as upper and low thresholds instead of calculating our own standard deviations and mean in the subsample with imaging data, because we have a much smaller sample, which may introduce more noise and exclude an excessive amount of people.

For smoking status, we used the self-reported smoking status information (<http://biobank.ctsu.ox.ac.uk/crystal/field.cgi?id=20116>). Participants could choose from one of the four options: (a) current smoker, (b) previous smoker, (c) non-smoker and (d) prefer not to answer. There were NAs for those did not answer. As the number of people who chose ‘prefer not to answer’ was very small, we did not transfer this into NA so to maximize our sample size. We treated this covariate as a categorical variable in our model. For the sensitivity analysis shown in Table S1, S4 and S5, in order to make it easier for demonstration, we transferred it into a numeric variable (current smoker = 2, previous smoker =1, non-smoker = 0, prefer not to answer = NA). This still represents the effects of smoking to depressive symptoms and white matter microstructure, as it is generally believed that there should be a gradient effect from current smoker to non-smoker.

Stressful life events described the number of events happened within 2 years before scanning session (<http://biobank.ctsu.ox.ac.uk/crystal/field.cgi?id=6145>). Items include: serious illness, injury or assault to oneself, death of a close relative, death of a spouse or partner, marital separation/divorce and financial difficulties.

**Other behavioural measures included in the present study**

All the behavioural variables were collected along with the imaging assessment unless it was collected within other categories such as online-follow up questionnaires that were not collected at any of the UK Biobank assessment centres.

Variables collected with the imaging assessment are listed below (relevant URL in the data showcase website shown in the brackets). For all the answers for the items listed below, “Prefer not to answer” and “Do not know” were recoded as NA.

1. Insomnia (<http://biobank.ctsu.ox.ac.uk/showcase/field.cgi?id=1200>), as indicated by 1=Never/rarely, 2=Sometimes, and 3=Usually.
2. Smoking status (<http://biobank.ctsu.ox.ac.uk/showcase/field.cgi?id=20116>), with 0=Never, 1=Previous, and 2=Current.
3. Recent pains in last month (<http://biobank.ctsu.ox.ac.uk/showcase/field.cgi?id=6159>), with 1=any type of pain reported and 0=None of the above. This variable was set as binary and logistic regression was used for its association tests. Therefore log odds ratios were reported (see Table 1).
4. Hand grip strength was derived as the mean of left- and right-hand grip strength (<http://biobank.ctsu.ox.ac.uk/showcase/field.cgi?id=46>, <http://biobank.ctsu.ox.ac.uk/showcase/field.cgi?id=47>).
5. g.Cognition. A selection of variables for cognitive functions was included to generate a g factor that represents the variance of general cognitive ability. Cognitive tasks include: Reaction time (<http://biobank.ctsu.ox.ac.uk/showcase/field.cgi?id=20023>), Verbal-numeric reasoning (also referred to as fluid intelligence in UK Biobank data dictionary: <http://biobank.ctsu.ox.ac.uk/showcase/field.cgi?id=20016>), Trail making (derived by Trail B – Trail A: <http://biobank.ctsu.ox.ac.uk/showcase/field.cgi?id=6350>, <http://biobank.ctsu.ox.ac.uk/showcase/field.cgi?id=6348>), Pair matching (<http://biobank.ctsu.ox.ac.uk/showcase/field.cgi?id=399>), Digit symbol substitution (<http://biobank.ctsu.ox.ac.uk/showcase/field.cgi?id=23324>), Matrix pattern completion (<http://biobank.ctsu.ox.ac.uk/showcase/field.cgi?id=6373>) and Tower rearranging (<http://biobank.ctsu.ox.ac.uk/showcase/field.cgi?id=21004>). A principal component analysis (PCA) was performed and the score of the first unrotated principal component was extracted as a measure of g factor for cognitive functions. The first component explained 34.5% of the total variance. Absolute correlation loadings for each item ranged from 0.350 to 0.691 (loadings for reaction time and pair matching were negative, as for these items a smaller reaction time is better).
6. g.Cognition for processing speed. An independent PCA was conducted on a selected range of the above tasks that has explicated stated measuring processing speed. The tasks included were: Reaction time, Trail making which tests visual processing speed, Digit symbol substitution which tests complex processing speed, and Pair matching, Matrix pattern completion, Tower rearranging and Verbal-numeric reasoning as participants were instructed to give answers in a limited time frame. Similar as g.Cognition, these measures used the score of the first unrotated principal component. The first principal component explained 42.2% of total variance. Absolute correlation loadings ranged from 0.553 to 0.734 (loading for reaction time was negative and all the others positive).
7. Neuroticism. Fields from 1920 to 2030 in the URL <http://biobank.ctsu.ox.ac.uk/showcase/label.cgi?id=100060> were used, and a total number of items with a ‘Yes’ answer for each participant was calculated as the neuroticism score.

Other variables that were not acquired with the imaging assessment were also often considered as lifetime variables that largely remain stable over time. These include the age of onset for depression (<http://biobank.ctsu.ox.ac.uk/showcase/field.cgi?id=20433>), household income (<http://biobank.ctsu.ox.ac.uk/showcase/field.cgi?id=738>), educational attainment (<http://biobank.ctsu.ox.ac.uk/showcase/field.cgi?id=6138>, 1=College or University degree, 0=others that are less than College or University degree), Townsend Index (<http://biobank.ctsu.ox.ac.uk/showcase/field.cgi?id=189>, 1=the least deprived tertile, 2=the second tertile, 3=the most deprived tertile).

**Supplementary results**

**Associations between measures of depressive symptoms and NODDI measures**

In general, results for ISOVF showed the largest resemblance with results for MD (see Figure S5). Here we report the results for all NODDI measures for each measure of depressive symptoms.

**Cross-sectional assessment of depressive symptoms**

For general variances of NODDI measures, higher cross-sectional depressive symptoms were associated with higher global ISOVF, higher ISOVF in gAF, gTR and gPF (β ranged from 0.018 to 0.029, p_corr_ ranged from 0.012 to 1.83×10^-5^). Higher cross-sectional measure was also associated with lower OD in gTR (β = -0.016, p_corr_ = 0.047).

Tract-specific associations were mainly found in ISOVF. Higher cross-sectional depressive symptoms was associated with higher ISOVF in anterior thalamic radiations, cingulate gyrus part of cingulum, parahippocampal part of cingulum, corticospinal tract, inferior fronto-occipital fasciculus, inferior longitudinal fasciculus, superior longitudinal fasciculus, superior thalamic radiation, uncinate fasciculus and middle cerebellar peduncle (β ranged from 0.015 to 0.039, p_corr_ ranged from 0.048 to 6.45×10^-8^). For other NODDI measures, higher cross-sectional measure was positively associated with OD in medial lemniscus (β = 0.018, p_corr_ = 0.050) and negatively associated with OD in superior thalamic radiation (β = -0.018, p_corr_ = 0.050).

No general or tract-specific associations were found for ICVF (p_corr_>0.314).

**Longitudinal trajectory of depressive symptoms**

For general variance in NODDI measures, a higher slope of longitudinal increase of depressive symptoms was associated with lower OD in global variance and gPF (β ranged from-0.035 to -0.039, p_corr_ ranged from 0.046 to 0.045). No general variance in ICVF showed a significant association with longitudinal slope (p_corr_>0.580).

For specific tracts, a steeper slope of longitudinal trajectory was associated with higher ISOVF in corticospinal tract and superior thalamic radiation (β ranged from 0.029 to 0.035, p_corr_ ranged from 0.040 to 0.025), as well as lower OD in inferior longitudinal fasciculus (β = -0.051, p_corr_ = 0.004). No tract measure was found associated with the slope of the longitudinal growth curve for ICVF measures (all p_corr_ > 0.314).

**The mean of depressive symptoms derived from multiple assessments**

Associations regarding mean depressive symptoms were mainly shown in ISOVF. Higher mean depressive symptoms were associated with higher ISOVF in global variance, gAF, gTR and gPF (β ranged from 0.014 to 0.027, p_corr_ ranged from 0.046 to 3.29×10^-4^). It is also associated with higher gAF OD (β = 0.016, p_corr_ = 0.047) and lower gTR OD (β = -0.016, p_corr_ = 0.049). Tract-wise analysis showed that ISOVF of anterior thalamic radiations, cingulate gyrus part of cingulum, corticospinal tract, superior longitudinal fasciculus, superior thalamic radiation, uncinate fasciculus and middle cerebellar peduncle (β ranged from 0.017 to 0.029, p_corr_ ranged from 0.021 to 5.32×10^-9^) were associated with higher mean depressive symptoms. Other than the associations with mean depressive symptoms shown in ISOVF, other association was found in lower OD in inferior longitudinal fasciculus (β = -0.051, p_corr_ = 4.04×10^-3^).

**The variability of depressive symptoms derived from multiple assessments**

For general variance, positive associations were shown in all g measures including global variance, gAF, gTR and gPF in ISOVF (β ranged from 0.016 to 0.033, p_corr_ ranged from 0.042 to 1.83×10^-4^). Negative associations in OD were found in global variance and gAF (β ranged from 0.019 to 0.023, p_corr_ ranged from 0.046 to 0.040).

Tract-wise analysis showed that variability of depressive symptoms were positively associated with ISOVF in anterior thalamic radiations, superior thalamic radiation, uncinate fasciculus and middle cerebellar peduncle (β ranged from 0.015 to 0.037, p_corr_ ranged from 0.048 to 5.64×10^-6^). Other than measures of ISOVF, other association was found between variability of depressive symptoms and OD in parahippocampal part of cingulum and medial lemniscus (β ranged from 0.020 to 0.021, p_corr_ ranged from 0.047 to 0.050).

For the above associations, we ran additional models to test if the results would change when smoking status, SLE and alcohol consumption were not corrected for. Results can be found in Figure S6. No association found in the main model where the three covariates were included turned insignificant after the covariates were removed.

**Effect of partial-volume contamination to FA and MD**

We observed differences of results for FA and MD. A possible explanation may be the different effect of partial volume contamination on FA and MD. To control for possible effects of partial-volume contamination related to structural atrophy, we have included age and age^2^ as covariates(3). We also conducted an additional analysis including brain size (in UK Biobank data dictionary: f.25010.2.0, described as “Volume of Brain, grey+white matter”) as one of the covariates and the results remained significant for both FA and MD except for one association turned null for MD in anterior thalamic radiation after controlling for brain size, however reached nominal significance before multiple correction applied (β = 0.030, p_uncorr_ = 0.020, p_corr_ = 0.053). (Figure S7).

Figure S1. Illustration of WM tracts. The tracts were defined by tractography mapping on FA (fractional anisotropy) data using AutoPtx (<https://fsl.fmrib.ox.ac.uk/fsl/fslwiki/AutoPtx>). They were categorised into three subsets as shown in the figure. Forceps major, forceps minor and uncinate fasciculus are unilateral structures and the rest are bilateral. For the purpose of clear illustration, bilateral structures were shown identically in both hemispheres.

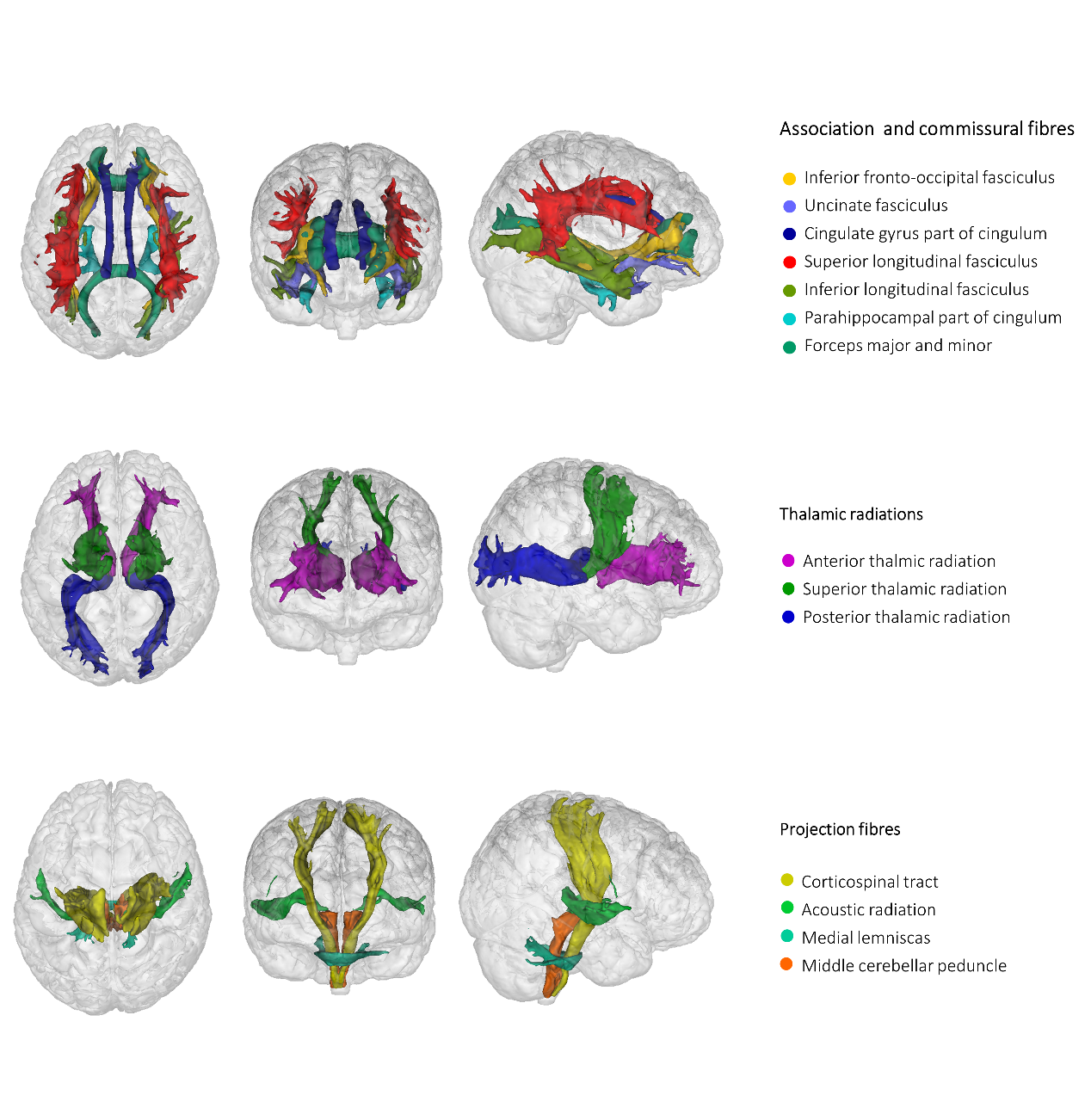

Figure S2. Description of sample sizes and changes due to each step of data merging or outlier removal. Cross-sectional = Cross-sectional measure at the imaging assessment, Mean = mean level of depressive symptoms based on multiple assessments for at least two times, Variability = standard deviation of depressive level of multiple assessments for at least three times, and Longitudinal slope = slope of longitudinal changes over all four times of assessments.

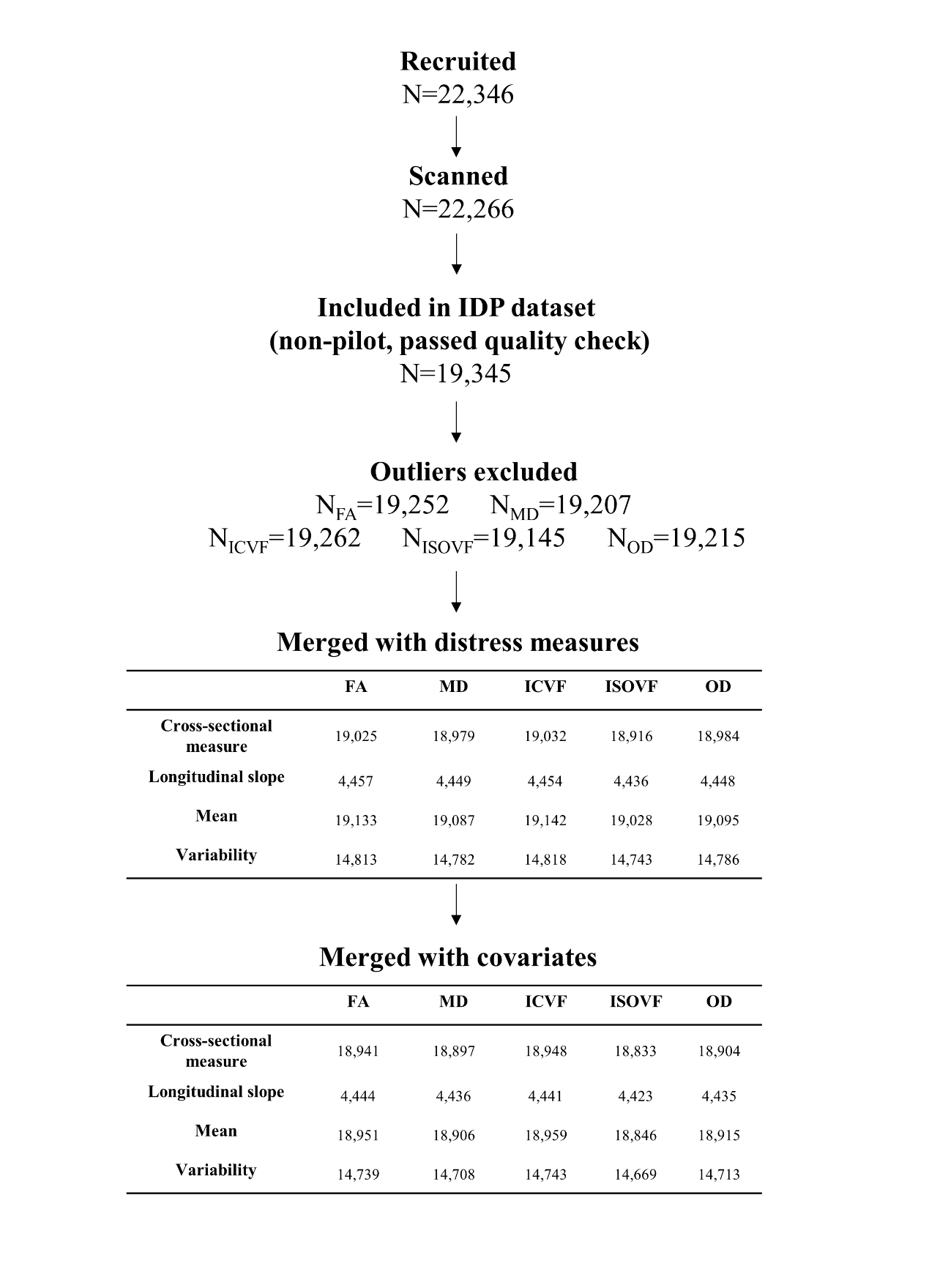

Figure S3. Distributions PHQ-4 and derived measures for depressive symptoms. (a) Density map for all four time points of assessments for PHQ-4. Instance 2 was for imaging assessment, and the depressive level acquired from this instance was used as a baseline, one-time measure for depressive symptoms. The mean depressive level over a minimum of two time points of assessment was also presented in this sub-figure (Depre.mean). (b) Density map for instability of depressive symptoms (Depre.variability). (c) Density map for the distribution of slope for the longitudinal trajectory of depressive level over four instances (Depre.longitudinal).

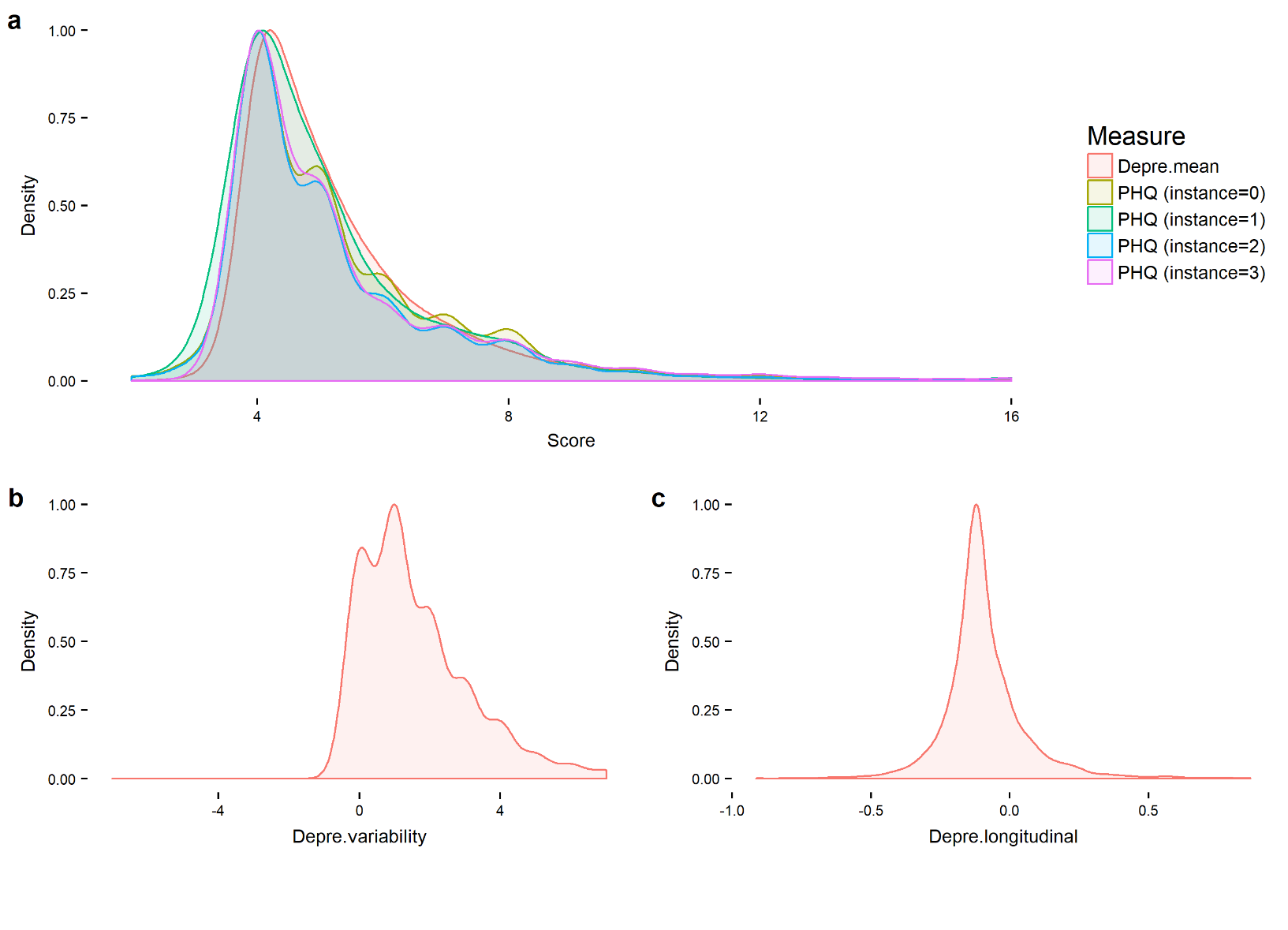

Figure S4. Variance explained by the principal components of PCA on total and subsets of white matter tracts.

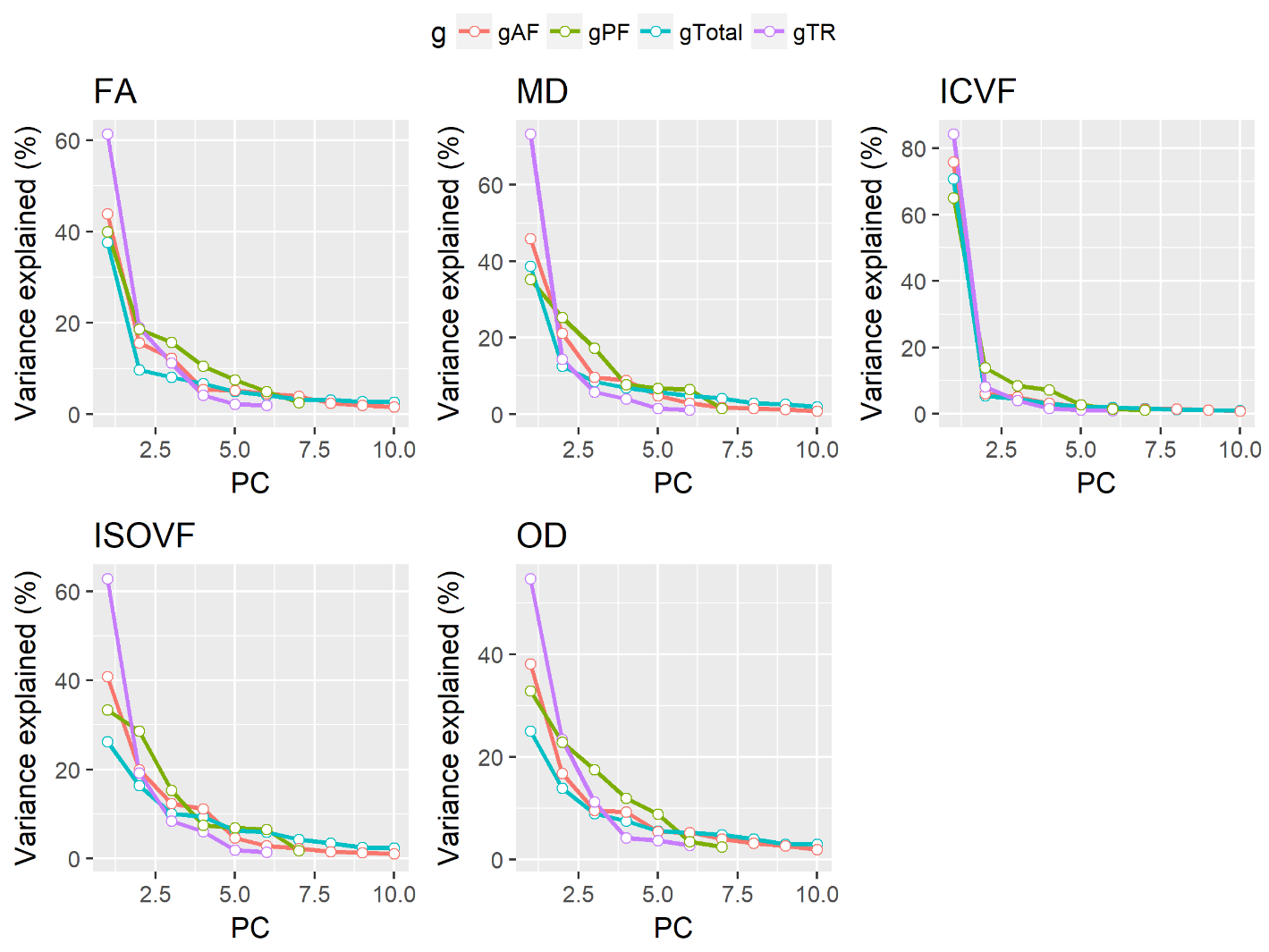

Figure S5. Results for all dMRI measures including DTI and NODDI. More descriptive statistics were shown in Table S2 and S3, results in the main text and supplementary results. In the heatmaps, each colour theme represents one dMRI measure. For the measures of depressive symptoms: Cross-sectional = depressive level at the imaging assessment, Longitudinal slope = slope of longitudinal changes over three time points until the imaging assessment, Mean = mean level of depressive symptoms based on multiple assessments for at least two times, Variability = standard deviation of depressive level of multiple assessments for at least three times. As the measures of FA, ICVF and OD have negative direction with MD and ISOVF, here in this figure, the effect sizes for FA, ICVF and OD were reversed (×-1). Significant associations after FDR correction were marked with a single asterisk.

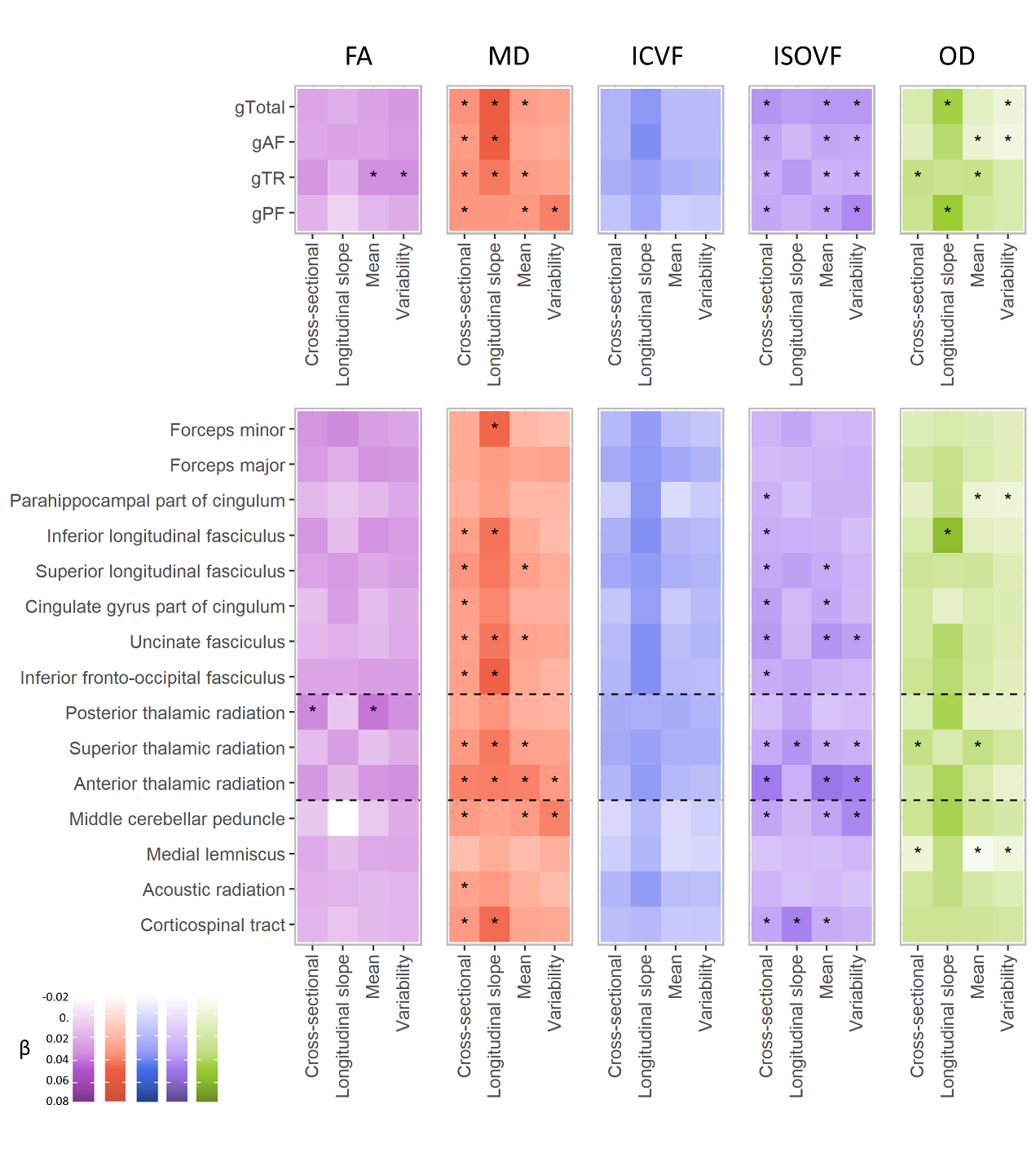

Figure S6. Results for a secondary model without controlling for stressful life events, smoking status or alcohol consumption. For the abbreviations and multiple correction methods, see the legend of Figure S5.

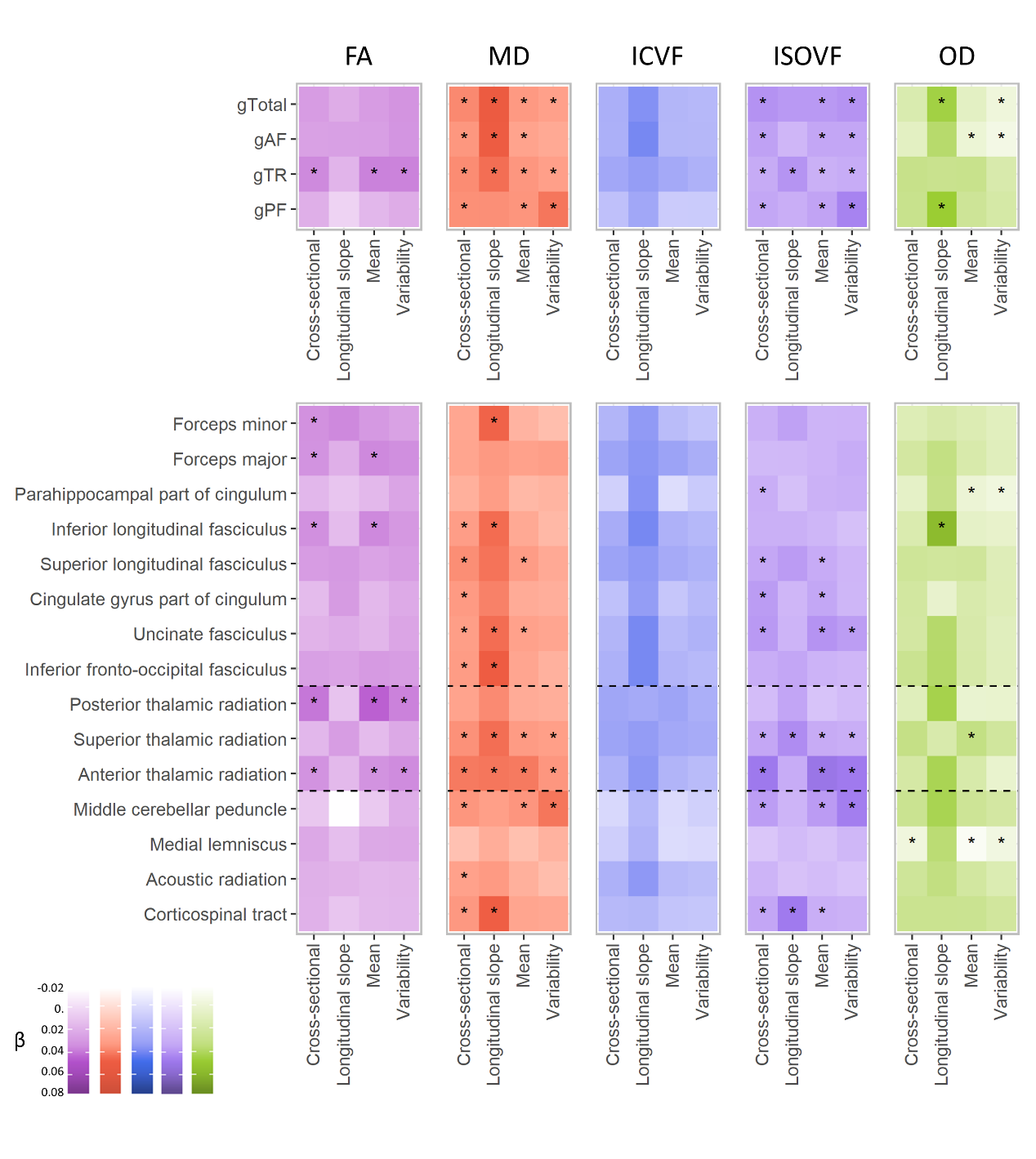

Figure S7. Results for adding brain size as a covariate.

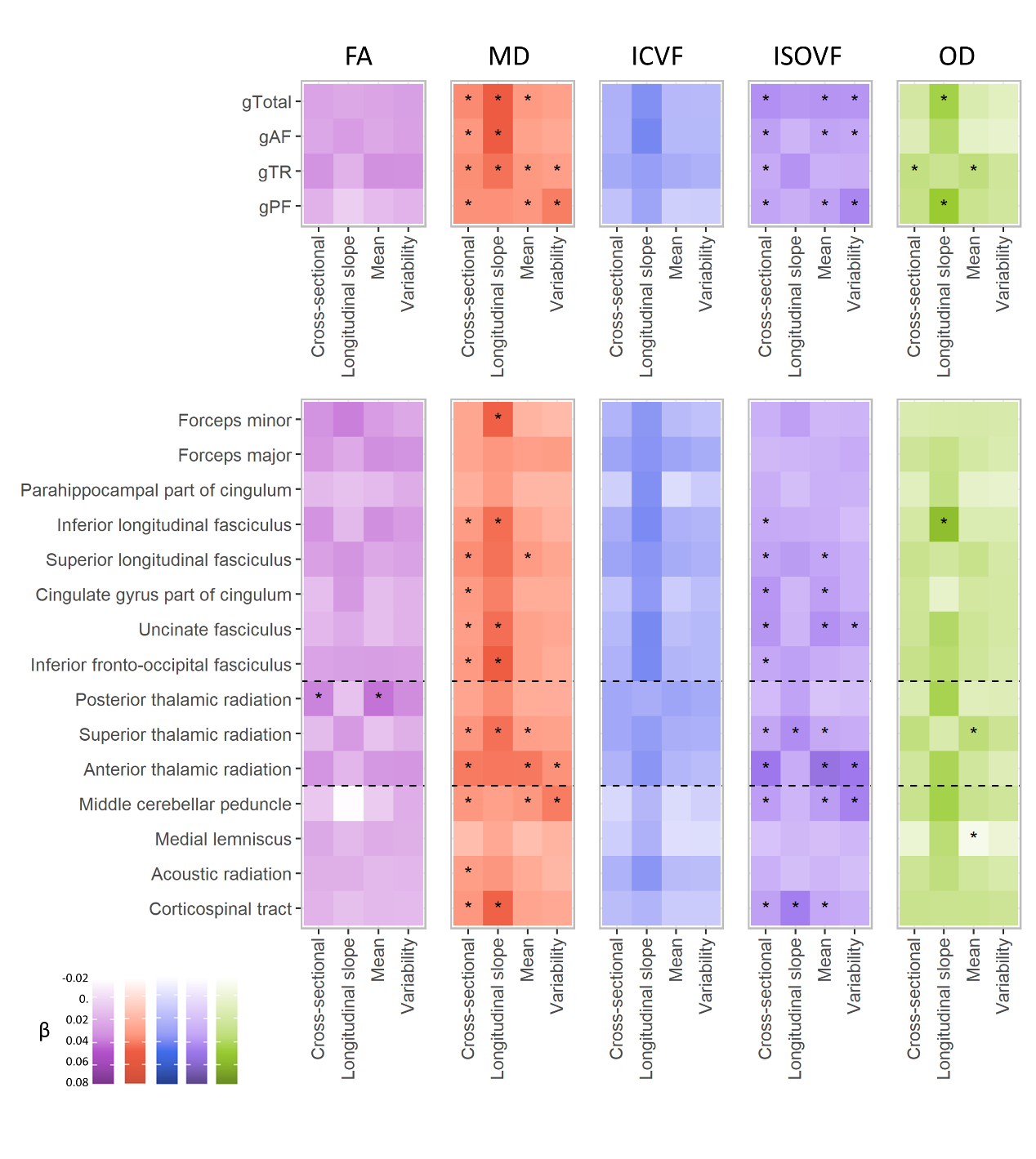

Table S1. Correlation matrix for all measures for depressive symptoms, smoking status, alcohol consumption, stressful life events, neuroticism and age range (time lag) for the multiple assessments used for generating mean level of depressive symptoms. For the measures of depressive symptoms: Cross-sectional measure = depressive level at the imaging assessment, Mean = mean level of depressive symptoms based on multiple assessments for at least two times, Variability = standard deviation of depressive level of multiple assessments for at least three times, and Longitudinal slope = slope of longitudinal changes over all four times of assessments. All r>0.3 were highlighted in bold. Measures for depressive symptoms were correlated with one another.

|  | Cross-sectional measure | Mean | Variability | Longitudinal slope | Smoking | Alcohol | SLE | N of occasions | Time.lag |
| --- | --- | --- | --- | --- | --- | --- | --- | --- | --- |
| Cross-sectional measure | 1 | **0.84** | **0.478** | **0.457** | 0.04 | -0.035 | 0.18 | -- | -- |
| Mean | **0.84** | 1 | **0.651** | -0.046 | 0.055 | -0.037 | 0.189 | -0.053 | 0.013 |
| Variability | **0.478** | **0.651** | 1 | -0.124 | 0.037 | -0.024 | 0.127 | 0.03 | 0.014 |
| Longitudinal slope | **0.457** | -0.046 | -0.124 | 1 | -0.002 | 0.005 | 0.041 | -- | 0.028 |
| Smoking | 0.04 | 0.055 | 0.037 | -0.002 | 1 | 0.2 | 0.005 | -0.013 | -0.011 |
| Alcohol | -0.035 | -0.037 | -0.024 | 0.005 | 0.2 | 1 | -0.036 | 0.001 | 0 |
| SLE | 0.18 | 0.189 | 0.127 | 0.041 | 0.005 | -0.036 | 1 | -0.002 | -0.008 |
| N of occasions | -- | -0.053 | 0.03 | -- | -0.013 | 0.001 | -0.002 | 1 | -- |
| Time.lag | -- | 0.013 | 0.014 | 0.028 | -0.011 | 0 | -0.008 | -- | 1 |

Table S2. Correlation loadings of each tract of PCA. For each dMRI measure, PCA on all tracts, association fibres, thalamic radiations and projection fibres were performed respectively. The loadings were reported as correlation loadings.

| **Tract** | **FA** | | | | **MD** | | | | **ICVF** | | | | **ISOVF** | | | | **OD** | | | |
| --- | --- | --- | --- | --- | --- | --- | --- | --- | --- | --- | --- | --- | --- | --- | --- | --- | --- | --- | --- | --- |
|  | **gTotal** | **gAF** | **gTR** | **gPF** | **gTotal** | **gAF** | **gTR** | **gPF** | **gTotal** | **gAF** | **gTR** | **gPF** | **gTotal** | **gAF** | **gTR** | **gPF** | **gTotal** | **gAF** | **gTR** | **gPF** |
| Parahippocampal part of cingulum | 0.361 | 0.379 | -- | -- | 0.472 | 0.61 | -- | -- | 0.604 | 0.61 | -- | -- | 0.63 | 0.845 | -- | -- | 0.791 | 0.88 | -- | -- |
| Parahippocampal part of cingulum | 0.394 | 0.419 | -- | -- | 0.454 | 0.587 | -- | -- | 0.627 | 0.636 | -- | -- | 0.652 | 0.865 | -- | -- | 0.737 | 0.846 | -- | -- |
| Forceps major | 0.547 | 0.552 | -- | -- | 0.487 | 0.518 | -- | -- | 0.796 | 0.793 | -- | -- | 0.187 | 0.122 | -- | -- | 0.192 | 0.127 | -- | -- |
| Cingulate gyrus part of cingulum | 0.558 | 0.635 | -- | -- | 0.61 | 0.576 | -- | -- | 0.783 | 0.801 | -- | -- | 0.346 | 0.191 | -- | -- | 0.216 | 0.17 | -- | -- |
| Cingulate gyrus part of cingulum | 0.583 | 0.658 | -- | -- | 0.615 | 0.59 | -- | -- | 0.787 | 0.805 | -- | -- | 0.331 | 0.205 | -- | -- | 0.201 | 0.149 | -- | -- |
| Uncinate fasciculus | 0.649 | 0.666 | -- | -- | 0.665 | 0.7 | -- | -- | 0.808 | 0.823 | -- | -- | 0.422 | 0.337 | -- | -- | 0.483 | 0.438 | -- | -- |
| Uncinate fasciculus | 0.681 | 0.696 | -- | -- | 0.734 | 0.75 | -- | -- | 0.841 | 0.856 | -- | -- | 0.493 | 0.356 | -- | -- | 0.523 | 0.481 | -- | -- |
| Superior longitudinal fasciculus | 0.81 | 0.795 | -- | -- | 0.859 | 0.795 | -- | -- | 0.931 | 0.933 | -- | -- | 0.608 | 0.3 | -- | -- | 0.317 | 0.125 | -- | -- |
| Superior longitudinal fasciculus | 0.824 | 0.8 | -- | -- | 0.847 | 0.777 | -- | -- | 0.928 | 0.929 | -- | -- | 0.622 | 0.292 | -- | -- | 0.356 | 0.153 | -- | -- |
| Forceps minor | 0.803 | 0.804 | -- | -- | 0.715 | 0.655 | -- | -- | 0.901 | 0.919 | -- | -- | 0.477 | 0.211 | -- | -- | 0.329 | 0.22 | -- | -- |
| Inferior longitudinal fasciculus | 0.822 | 0.813 | -- | -- | 0.862 | 0.837 | -- | -- | 0.939 | 0.941 | -- | -- | 0.617 | 0.319 | -- | -- | 0.492 | 0.409 | -- | -- |
| Inferior longitudinal fasciculus | 0.85 | 0.827 | -- | -- | 0.893 | 0.85 | -- | -- | 0.943 | 0.94 | -- | -- | 0.663 | 0.333 | -- | -- | 0.56 | 0.473 | -- | -- |
| Inferior fronto-occipital fasciculus | 0.827 | 0.83 | -- | -- | 0.871 | 0.836 | -- | -- | 0.946 | 0.951 | -- | -- | 0.624 | 0.344 | -- | -- | 0.461 | 0.383 | -- | -- |
| Inferior fronto-occipital fasciculus | 0.853 | 0.842 | -- | -- | 0.89 | 0.849 | -- | -- | 0.951 | 0.955 | -- | -- | 0.633 | 0.332 | -- | -- | 0.492 | 0.398 | -- | -- |
| Superior thalamic radiation | 0.654 | -- | 0.731 | -- | 0.773 | -- | 0.844 | -- | 0.882 | -- | 0.921 | -- | 0.589 | -- | 0.694 | -- | 0.425 | -- | 0.907 | -- |
| Superior thalamic radiation | 0.65 | -- | 0.737 | -- | 0.742 | -- | 0.82 | -- | 0.874 | -- | 0.916 | -- | 0.562 | -- | 0.674 | -- | 0.445 | -- | 0.915 | -- |
| Posterior thalamic radiation | 0.675 | -- | 0.785 | -- | 0.756 | -- | 0.863 | -- | 0.872 | -- | 0.902 | -- | 0.626 | -- | 0.879 | -- | 0.436 | -- | 0.271 | -- |
| Anterior thalamic radiation | 0.774 | -- | 0.807 | -- | 0.82 | -- | 0.85 | -- | 0.902 | -- | 0.933 | -- | 0.573 | -- | 0.585 | -- | 0.495 | -- | 0.558 | -- |
| Posterior thalamic radiation | 0.693 | -- | 0.812 | -- | 0.774 | -- | 0.88 | -- | 0.879 | -- | 0.906 | -- | 0.656 | -- | 0.897 | -- | 0.456 | -- | 0.33 | -- |
| Anterior thalamic radiation | 0.773 | -- | 0.816 | -- | 0.811 | -- | 0.847 | -- | 0.895 | -- | 0.931 | -- | 0.572 | -- | 0.597 | -- | 0.503 | -- | 0.655 | -- |
| Medial lemniscus | 0.262 | -- | -- | 0.501 | 0.253 | -- | -- | -0.328 | 0.492 | -- | -- | 0.737 | 0.42 | -- | -- | -0.645 | 0.169 | -- | -- | 0.22 |
| Medial lemniscus | 0.273 | -- | -- | 0.519 | 0.222 | -- | -- | -0.309 | 0.48 | -- | -- | 0.733 | 0.379 | -- | -- | -0.63 | 0.278 | -- | -- | 0.258 |
| Middle cerebellar peduncle | 0.365 | -- | -- | 0.539 | 0.35 | -- | -- | -0.933 | 0.503 | -- | -- | 0.736 | 0.355 | -- | -- | -0.753 | 0.354 | -- | -- | 0.862 |
| Acoustic radiation | 0.608 | -- | -- | 0.577 | 0.487 | -- | -- | -0.305 | 0.875 | -- | -- | 0.861 | 0.267 | -- | -- | -0.173 | 0.255 | -- | -- | 0.312 |
| Acoustic radiation | 0.618 | -- | -- | 0.631 | 0.505 | -- | -- | -0.274 | 0.854 | -- | -- | 0.854 | 0.292 | -- | -- | -0.151 | 0.351 | -- | -- | 0.345 |
| Corticospinal tract | 0.576 | -- | -- | 0.802 | 0.579 | -- | -- | -0.282 | 0.789 | -- | -- | 0.85 | 0.319 | -- | -- | -0.152 | 0.34 | -- | -- | 0.552 |
| Corticospinal tract | 0.581 | -- | -- | 0.821 | 0.58 | -- | -- | -0.295 | 0.778 | -- | -- | 0.844 | 0.36 | -- | -- | -0.21 | 0.402 | -- | -- | 0.585 |

Table S3. Main results for DTI measures (FA and MD). Betas were standardised effect sizes. P values were un-corrected p values. All p_corrected_<0.05 were marked by asterixis. FDR correction was applied on all association tests for a dMRI measure (n = 15*4 measures of depressive symptoms = 60 for tract analysis, and n = 4*4 measures of depressive symptoms = 16 for g analysis). For the measures of depressive symptoms: Cross-sectional measure = depressive level at the imaging assessment, Mean = mean level of depressive symptoms based on multiple assessments for at least two times, Variability = standard deviation of depressive level of multiple assessments for at least three times, and Longitudinal slope = slope of longitudinal changes over all four times of assessments.

|  |  | **Cross-sectional measure** | | **Mean** | | **Variability** | | **Longitudinal slope** | |
| --- | --- | --- | --- | --- | --- | --- | --- | --- | --- |
| **Tract name** | **Measure** | **Beta (Std)** | **p** | **Beta (Std)** | **p** | **Beta (Std)** | **p** | **Beta (Std)** | **p** |
| g.Total | FA | -0.011 (0.007) | 0.116 | -0.012 (0.007) | 0.099 | -0.017 (0.008) | 0.036 | -0.006 (0.014) | 0.679 |
|  | MD | 0.024 (0.007) | 4.17e-04* | 0.018 (0.007) | 0.008* | 0.015 (0.007) | 0.052 | 0.04 (0.014) | 0.003* |
| g.AF | FA | -0.009 (0.007) | 0.196 | -0.01 (0.007) | 0.145 | -0.016 (0.008) | 0.047 | -0.012 (0.014) | 0.397 |
|  | MD | 0.019 (0.007) | 0.005* | 0.014 (0.007) | 0.048 | 0.01 (0.008) | 0.192 | 0.039 (0.014) | 0.005* |
| g.TR | FA | -0.019 (0.007) | 0.01 | -0.021 (0.007) | 0.004* | -0.022 (0.008) | 0.006* | -0.001 (0.015) | 0.927 |
|  | MD | 0.022 (0.006) | 4.37e-04* | 0.018 (0.006) | 0.004* | 0.015 (0.007) | 0.039 | 0.031 (0.013) | 0.017* |
| g.PF | FA | -0.004 (0.007) | 0.611 | -9.55e-04 (0.007) | 0.895 | -0.007 (0.008) | 0.371 | 0.013 (0.014) | 0.35 |
|  | MD | 0.022 (0.007) | 0.002* | 0.02 (0.007) | 0.004* | 0.029 (0.008) | 2.29e-04* | 0.022 (0.014) | 0.119 |
| Acoustic radiation | FA | -0.004 (0.006) | 0.564 | 1.60e-04 (0.006) | 0.98 | -0.002 (0.007) | 0.787 | -0.003 (0.013) | 0.785 |
|  | MD | 0.015 (0.006) | 0.014* | 0.008 (0.006) | 0.222 | 5.81e-04 (0.007) | 0.933 | 0.02 (0.013) | 0.113 |
| Anterior thalamic radiation | FA | -0.018 (0.007) | 0.009 | -0.018 (0.007) | 0.008 | -0.021 (0.008) | 0.006 | 5.61e-04 (0.014) | 0.967 |
|  | MD | 0.028 (0.006) | 4.88e-06* | 0.029 (0.006) | 4.20e-06* | 0.02 (0.007) | 0.003* | 0.03 (0.013) | 0.02* |
| Cingulate gyrus part of cingulum | FA | 0.003 (0.006) | 0.638 | 0.001 (0.006) | 0.876 | -0.006 (0.007) | 0.378 | -0.014 (0.013) | 0.271 |
|  | MD | 0.017 (0.007) | 0.01* | 0.008 (0.007) | 0.241 | 0.007 (0.007) | 0.294 | 0.027 (0.013) | 0.037 |
| Parahippocampal part of cingulum | FA | -1.10e-04 (0.006) | 0.986 | -6.36e-04 (0.006) | 0.921 | -0.01 (0.007) | 0.175 | 0.006 (0.013) | 0.621 |
|  | MD | 0.007 (0.006) | 0.256 | 0.004 (0.006) | 0.554 | 0.004 (0.007) | 0.546 | 0.017 (0.012) | 0.17 |
| Corticospinal tract | FA | -0.003 (0.007) | 0.699 | -7.23e-04 (0.007) | 0.915 | -0.002 (0.008) | 0.838 | 0.006 (0.014) | 0.679 |
|  | MD | 0.02 (0.007) | 0.003* | 0.014 (0.007) | 0.046 | 0.012 (0.007) | 0.11 | 0.035 (0.013) | 0.007* |
| Inferior fronto occipital fasciculus | FA | -0.01 (0.007) | 0.133 | -0.014 (0.007) | 0.053 | -0.013 (0.008) | 0.085 | -0.011 (0.014) | 0.451 |
|  | MD | 0.017 (0.007) | 0.011* | 0.012 (0.007) | 0.064 | 0.006 (0.007) | 0.412 | 0.038 (0.013) | 0.004* |
| Inferior longitudinal fasciculus | FA | -0.018 (0.007) | 0.009 | -0.02 (0.007) | 0.003 | -0.015 (0.008) | 0.052 | 0.002 (0.014) | 0.911 |
|  | MD | 0.016 (0.007) | 0.013* | 0.01 (0.007) | 0.117 | 0.002 (0.007) | 0.755 | 0.033 (0.013) | 0.013* |
| Medial lemniscus | FA | -0.008 (0.006) | 0.204 | -0.008 (0.006) | 0.201 | -0.009 (0.007) | 0.194 | 0.003 (0.012) | 0.833 |
|  | MD | -1.75e-04 (0.006) | 0.978 | 8.64e-04 (0.006) | 0.891 | 0.008 (0.007) | 0.229 | 0.009 (0.012) | 0.465 |
| Posterior thalamic radiation | FA | -0.024 (0.007) | 3.75e-04* | -0.029 (0.007) | 1.31e-05* | -0.022 (0.007) | 0.003 | 0.007 (0.014) | 0.62 |
|  | MD | 0.012 (0.006) | 0.049 | 0.007 (0.006) | 0.256 | 0.006 (0.007) | 0.357 | 0.023 (0.013) | 0.075 |
| Superior longitudinal fasciculus | FA | -0.011 (0.007) | 0.098 | -0.009 (0.007) | 0.201 | -0.013 (0.008) | 0.091 | -0.016 (0.014) | 0.24 |
|  | MD | 0.022 (0.007) | 0.001* | 0.017 (0.007) | 0.015* | 0.01 (0.008) | 0.18 | 0.031 (0.014) | 0.024 |
| Superior thalamic radiation | FA | 0.002 (0.007) | 0.794 | 0.004 (0.007) | 0.615 | -0.007 (0.008) | 0.369 | -0.014 (0.014) | 0.344 |
|  | MD | 0.02 (0.006) | 7.28e-04* | 0.016 (0.006) | 0.009* | 0.015 (0.007) | 0.026 | 0.031 (0.013) | 0.013* |
| Uncinate fasciculus | FA | -8.67e-04 (0.006) | 0.893 | 6.92e-04 (0.006) | 0.915 | -0.008 (0.007) | 0.281 | -0.005 (0.013) | 0.723 |
|  | MD | 0.016 (0.006) | 0.009* | 0.015 (0.006) | 0.017* | 0.013 (0.007) | 0.062 | 0.032 (0.012) | 0.01* |
| Forceps major | FA | -0.015 (0.007) | 0.038 | -0.019 (0.007) | 0.008 | -0.018 (0.008) | 0.029 | -0.006 (0.015) | 0.684 |
|  | MD | 0.011 (0.007) | 0.11 | 0.014 (0.007) | 0.049 | 0.015 (0.008) | 0.06 | 0.02 (0.014) | 0.166 |
| Forceps minor | FA | -0.019 (0.007) | 0.009 | -0.015 (0.007) | 0.037 | -0.011 (0.008) | 0.181 | -0.024 (0.014) | 0.102 |
|  | MD | 0.011 (0.007) | 0.114 | 0.003 (0.007) | 0.635 | -7.27e-04 (0.008) | 0.924 | 0.037 (0.014) | 0.009* |
| Middle cerebellar peduncle | FA | 0.008 (0.007) | 0.285 | 0.009 (0.007) | 0.238 | -0.007 (0.008) | 0.403 | 0.036 (0.015) | 0.014 |
|  | MD | 0.02 (0.007) | 0.006* | 0.019 (0.007) | 0.008* | 0.029 (0.008) | 3.87e-04* | 0.015 (0.014) | 0.301 |

Table S4. Main results for NODDI measures (ICVF, ISOVF, and OD). Betas were standardised effect sizes. For the abbreviations of measures for depressive symptoms, see the legend of Table S2.

|  |  | **Cross-sectional measure** | | **Mean** | | **Variability** | | **Longitudinal measure** | |
| --- | --- | --- | --- | --- | --- | --- | --- | --- | --- |
| **Tract name** | **Measure** | **Beta (Std)** | **p** | **Beta (Std)** | **p** | **Beta (Std)** | **p** | **Beta (Std)** | **p** |
| g.Total | ICVF | -0.007 (0.007) | 0.313 | -0.003 (0.007) | 0.636 | -0.003 (0.008) | 0.696 | -0.024 (0.014) | 0.094 |
|  | ISOVF | 0.029 (0.007) | 1.49e-05* | 0.027 (0.007) | 6.16e-05* | 0.027 (0.007) | 2.54e-04* | 0.025 (0.014) | 0.062 |
|  | OD | -0.001 (0.007) | 0.86 | 0.008 (0.007) | 0.233 | 0.019 (0.008) | 0.011* | -0.035 (0.014) | 0.011* |
| g.AF | ICVF | -0.007 (0.007) | 0.337 | -0.003 (0.007) | 0.669 | -0.004 (0.008) | 0.65 | -0.028 (0.014) | 0.057 |
|  | ISOVF | 0.022 (0.007) | 0.002* | 0.022 (0.007) | 0.003* | 0.02 (0.008) | 0.012* | 0.01 (0.014) | 0.495 |
|  | OD | 0.007 (0.007) | 0.285 | 0.016 (0.007) | 0.018* | 0.023 (0.008) | 0.003* | -0.026 (0.014) | 0.056 |
| g.TR | ICVF | -0.012 (0.007) | 0.098 | -0.01 (0.007) | 0.151 | -0.007 (0.008) | 0.386 | -0.019 (0.014) | 0.191 |
|  | ISOVF | 0.018 (0.006) | 0.007* | 0.014 (0.007) | 0.034* | 0.016 (0.007) | 0.029* | 0.027 (0.013) | 0.042 |
|  | OD | -0.016 (0.007) | 0.015* | -0.016 (0.007) | 0.022* | -4.87e-04 (0.007) | 0.948 | -0.013 (0.014) | 0.326 |
| g.PF | ICVF | 0.002 (0.007) | 0.782 | 0.009 (0.007) | 0.218 | 0.007 (0.008) | 0.401 | -0.014 (0.014) | 0.328 |
|  | ISOVF | 0.02 (0.007) | 0.004* | 0.022 (0.007) | 0.002* | 0.033 (0.008) | 2.29e-05* | 0.014 (0.014) | 0.304 |
|  | OD | -0.014 (0.007) | 0.054 | -0.009 (0.007) | 0.216 | -0.001 (0.008) | 0.899 | -0.039 (0.014) | 0.006* |
| Acoustic radiation | ICVF | -0.007 (0.007) | 0.315 | -0.002 (0.007) | 0.797 | -7.31e-04 (0.008) | 0.925 | -0.023 (0.014) | 0.11 |
|  | ISOVF | 0.013 (0.006) | 0.041 | 0.009 (0.006) | 0.173 | 0.002 (0.007) | 0.739 | 0.004 (0.013) | 0.727 |
|  | OD | -0.008 (0.006) | 0.185 | -0.004 (0.006) | 0.461 | 0.004 (0.007) | 0.582 | -0.02 (0.012) | 0.095 |
| Anterior thalamic radiation | ICVF | -0.007 (0.007) | 0.324 | -0.005 (0.007) | 0.445 | -0.002 (0.008) | 0.789 | -0.023 (0.014) | 0.101 |
|  | ISOVF | 0.039 (0.006) | 2.15e-09* | 0.042 (0.007) | 8.86e-11* | 0.037 (0.007) | 2.82e-07* | 0.015 (0.013) | 0.232 |
|  | OD | -0.004 (0.007) | 0.558 | -0.002 (0.007) | 0.819 | 0.014 (0.008) | 0.066 | -0.031 (0.014) | 0.024 |
| Cingulate gyrus part of cingulum | ICVF | 0.003 (0.007) | 0.691 | 0.007 (0.007) | 0.343 | -0.003 (0.008) | 0.711 | -0.021 (0.014) | 0.134 |
|  | ISOVF | 0.025 (0.006) | 6.68e-05* | 0.021 (0.006) | 0.001* | 0.01 (0.007) | 0.162 | 0.009 (0.013) | 0.464 |
|  | OD | -0.007 (0.006) | 0.276 | -8.22e-04 (0.006) | 0.894 | 0.003 (0.007) | 0.662 | 0.013 (0.012) | 0.304 |
| Parahippocampal part of cingulum | ICVF | 0.009 (0.007) | 0.156 | 0.016 (0.007) | 0.017 | 0.006 (0.007) | 0.424 | -0.024 (0.014) | 0.074 |
|  | ISOVF | 0.015 (0.006) | 0.018* | 0.014 (0.006) | 0.023 | 0.014 (0.007) | 0.039 | 0.004 (0.012) | 0.759 |
|  | OD | 0.01 (0.006) | 0.098 | 0.016 (0.006) | 0.006* | 0.02 (0.007) | 0.002* | -0.018 (0.012) | 0.146 |
| Corticospinal tract | ICVF | -0.001 (0.007) | 0.851 | 0.006 (0.007) | 0.43 | 0.005 (0.008) | 0.523 | -0.005 (0.014) | 0.725 |
|  | ISOVF | 0.022 (0.007) | 0.001* | 0.019 (0.007) | 0.005* | 0.014 (0.007) | 0.056 | 0.036 (0.013) | 0.008* |
|  | OD | -0.012 (0.007) | 0.06 | -0.01 (0.007) | 0.117 | -0.006 (0.007) | 0.415 | -0.012 (0.013) | 0.343 |
| Inferior fronto occipital fasciculus | ICVF | -0.006 (0.007) | 0.358 | -0.004 (0.007) | 0.554 | -0.002 (0.008) | 0.773 | -0.027 (0.014) | 0.059 |
|  | ISOVF | 0.018 (0.007) | 0.009* | 0.015 (0.007) | 0.032 | 0.011 (0.008) | 0.155 | 0.022 (0.014) | 0.102 |
|  | OD | -0.012 (0.007) | 0.06 | -0.003 (0.007) | 0.671 | 0.007 (0.007) | 0.336 | -0.025 (0.013) | 0.058 |
| Inferior longitudinal fasciculus | ICVF | -0.009 (0.007) | 0.202 | -0.006 (0.007) | 0.395 | -0.003 (0.008) | 0.701 | -0.027 (0.014) | 0.063 |
|  | ISOVF | 0.017 (0.007) | 0.013* | 0.013 (0.007) | 0.052 | 0.005 (0.008) | 0.482 | 0.015 (0.014) | 0.274 |
|  | OD | -5.82e-04 (0.006) | 0.928 | 0.009 (0.007) | 0.178 | 0.012 (0.007) | 0.082 | -0.051 (0.013) | 1.35e-04* |
| Medial lemniscus | ICVF | 0.008 (0.006) | 0.18 | 0.016 (0.006) | 0.007 | 0.013 (0.007) | 0.049 | -0.006 (0.012) | 0.609 |
|  | ISOVF | 0.002 (0.006) | 0.765 | 0.005 (0.006) | 0.426 | 0.011 (0.007) | 0.109 | 0.007 (0.012) | 0.551 |
|  | OD | 0.018 (0.007) | 0.006* | 0.03 (0.007) | 1.14e-05* | 0.021 (0.007) | 0.006* | -0.023 (0.013) | 0.087 |
| Posterior thalamic radiation | ICVF | -0.012 (0.007) | 0.079 | -0.012 (0.007) | 0.077 | -0.007 (0.008) | 0.334 | -0.01 (0.014) | 0.466 |
|  | ISOVF | 0.007 (0.006) | 0.285 | 0.002 (0.006) | 0.77 | 0.006 (0.007) | 0.376 | 0.021 (0.013) | 0.106 |
|  | OD | 0.004 (0.006) | 0.554 | 0.012 (0.006) | 0.053 | 0.013 (0.007) | 0.053 | -0.033 (0.012) | 0.008 |
| Superior longitudinal fasciculus | ICVF | -0.015 (0.007) | 0.035 | -0.011 (0.007) | 0.139 | -0.008 (0.008) | 0.339 | -0.023 (0.014) | 0.118 |
|  | ISOVF | 0.02 (0.007) | 0.003* | 0.018 (0.007) | 0.006* | 0.011 (0.007) | 0.15 | 0.024 (0.014) | 0.075 |
|  | OD | -0.011 (0.006) | 0.07 | -0.01 (0.006) | 0.133 | 0.005 (0.007) | 0.501 | -0.007 (0.013) | 0.555 |
| Superior thalamic radiation | ICVF | -0.013 (0.007) | 0.052 | -0.01 (0.007) | 0.137 | -0.01 (0.008) | 0.202 | -0.019 (0.014) | 0.178 |
|  | ISOVF | 0.019 (0.006) | 0.001* | 0.017 (0.006) | 0.003* | 0.015 (0.006) | 0.018* | 0.029 (0.012) | 0.014* |
|  | OD | -0.018 (0.007) | 0.007* | -0.019 (0.007) | 0.005* | -0.007 (0.007) | 0.338 | -1.92e-04 (0.013) | 0.988 |
| Uncinate fasciculus | ICVF | -0.004 (0.007) | 0.535 | -0.002 (0.007) | 0.817 | -0.006 (0.007) | 0.402 | -0.027 (0.013) | 0.047 |
|  | ISOVF | 0.027 (0.006) | 4.37e-05* | 0.029 (0.007) | 1.05e-05* | 0.024 (0.007) | 7.46e-04* | 0.01 (0.013) | 0.436 |
|  | OD | -0.007 (0.006) | 0.259 | -0.004 (0.006) | 0.494 | 0.004 (0.007) | 0.533 | -0.027 (0.012) | 0.028 |
| Forceps major | ICVF | -0.014 (0.007) | 0.059 | -0.014 (0.007) | 0.056 | -0.008 (0.008) | 0.332 | -0.023 (0.015) | 0.117 |
|  | ISOVF | 0.008 (0.007) | 0.293 | 0.012 (0.007) | 0.094 | 0.015 (0.008) | 0.055 | 0.01 (0.014) | 0.495 |
|  | OD | -0.007 (0.007) | 0.325 | -0.002 (0.007) | 0.817 | 0.006 (0.008) | 0.469 | -0.016 (0.014) | 0.262 |
| Forceps minor | ICVF | -0.005 (0.007) | 0.465 | -0.001 (0.007) | 0.871 | 0.003 (0.008) | 0.693 | -0.022 (0.014) | 0.122 |
|  | ISOVF | 0.013 (0.007) | 0.059 | 0.009 (0.007) | 0.184 | 0.011 (0.008) | 0.169 | 0.022 (0.014) | 0.109 |
|  | OD | 0.004 (0.007) | 0.591 | 0.003 (0.007) | 0.685 | 0.008 (0.008) | 0.343 | -0.002 (0.015) | 0.91 |
| Middle cerebellar peduncle | ICVF | 0.015 (0.007) | 0.037 | 0.016 (0.007) | 0.025 | 0.009 (0.008) | 0.254 | -0.004 (0.013) | 0.781 |
|  | ISOVF | 0.023 (0.007) | 0.002* | 0.023 (0.007) | 0.002* | 0.034 (0.008) | 3.58e-05* | 0.01 (0.014) | 0.485 |
|  | OD | -0.012 (0.007) | 0.098 | -0.01 (0.007) | 0.168 | -0.003 (0.008) | 0.669 | -0.033 (0.015) | 0.022 |

Table S5. The effects of stressful life events (SLE), neuroticism, smoking status, and alcohol consumption as covariates for DTI measures. The test was conducted on the full sample of IDP from UK Biobank imaging team, without outlier exclusion or phenotypic data merging. Betas were standardised effect sizes. P values were un-corrected p values. All p_corrected_<0.05 were marked by asterixis. FDR correction was applied on all tests conducted on one dMRI measure.

|  |  | **Alcohol consumption** | | **Smoking** | | **SLE** | |
| --- | --- | --- | --- | --- | --- | --- | --- |
| **Tract name** | **Measure** | **Beta (Std)** | **p** | **Beta (Std)** | **p** | **Beta (Std)** | **p** |
| g.Total | FA | -0.014 (0.007) | 0.053 | -0.016 (0.007) | 0.02* | -0.015 (0.007) | 0.031 |
|  | MD | 0.046 (0.007) | 1.04e-11* | 0.034 (0.007) | 2.31e-07* | 0.007 (0.007) | 0.316 |
| g.AF | FA | -0.023 (0.007) | 0.002* | -0.021 (0.007) | 0.003* | -0.014 (0.007) | 0.038 |
|  | MD | 0.034 (0.007) | 6.87e-07* | 0.022 (0.007) | 8.80e-04* | 0.006 (0.007) | 0.336 |
| g.TR | FA | -0.024 (0.007) | 0.001* | -0.031 (0.007) | 7.17e-06* | -0.015 (0.007) | 0.034 |
|  | MD | 0.057 (0.006) | 7.68e-19* | 0.049 (0.006) | 1.36e-15* | 0.006 (0.006) | 0.349 |
| g.PF | FA | 0.03 (0.007) | 2.13e-05* | 0.017 (0.007) | 0.012* | -0.009 (0.007) | 0.191 |
|  | MD | 0.05 (0.007) | 1.77e-12* | 0.052 (0.007) | 2.53e-14* | -2.58e-04 (0.007) | 0.97 |
| Acoustic radiation | FA | 0.009 (0.006) | 0.151 | 0.018 (0.006) | 0.003* | -0.01 (0.006) | 0.089 |
|  | MD | -2.94e-04 (0.006) | 0.963 | -0.01 (0.006) | 0.09 | 0.005 (0.006) | 0.386 |
| Anterior thalamic radiation | FA | -0.022 (0.007) | 0.001* | -0.009 (0.007) | 0.161 | -0.009 (0.007) | 0.155 |
|  | MD | 0.038 (0.006) | 1.69e-09* | 0.028 (0.006) | 2.94e-06* | 0.007 (0.006) | 0.272 |
| Cingulate gyrus part of cingulum | FA | -0.027 (0.006) | 2.23e-05* | -0.024 (0.006) | 9.46e-05* | -0.004 (0.006) | 0.481 |
|  | MD | 0.013 (0.007) | 0.049 | 0.011 (0.006) | 0.097 | 0.009 (0.006) | 0.15 |
| Parahippocampal part of cingulum | FA | 0.006 (0.006) | 0.381 | 6.89e-04 (0.006) | 0.911 | -0.002 (0.006) | 0.691 |
|  | MD | 0.012 (0.006) | 0.068 | 0.013 (0.006) | 0.038 | -0.001 (0.006) | 0.835 |
| Corticospinal tract | FA | 0.027 (0.007) | 8.18e-05* | 0.01 (0.007) | 0.112 | -0.004 (0.007) | 0.52 |
|  | MD | 0.021 (0.007) | 0.002* | 8.87e-04 (0.007) | 0.894 | 0.006 (0.007) | 0.392 |
| Inferior fronto occipital fasciculus | FA | -0.009 (0.007) | 0.182 | -0.013 (0.007) | 0.049 | -0.009 (0.007) | 0.187 |
|  | MD | 0.036 (0.007) | 8.81e-08* | 0.017 (0.006) | 0.008* | 0.005 (0.006) | 0.431 |
| Inferior longitudinal fasciculus | FA | -0.019 (0.007) | 0.005* | -0.024 (0.007) | 2.55e-04* | -0.011 (0.007) | 0.109 |
|  | MD | 0.039 (0.007) | 4.39e-09* | 0.021 (0.006) | 9.27e-04* | 0.004 (0.006) | 0.547 |
| Medial lemniscus | FA | 0.014 (0.006) | 0.022* | 0.002 (0.006) | 0.761 | -0.004 (0.006) | 0.548 |
|  | MD | 0.006 (0.006) | 0.309 | 0.001 (0.006) | 0.848 | -0.011 (0.006) | 0.08 |
| Posterior thalamic radiation | FA | -0.03 (0.007) | 8.07e-06* | -0.045 (0.006) | 4.79e-12* | -0.015 (0.007) | 0.023 |
|  | MD | 0.059 (0.006) | 1.65e-20* | 0.051 (0.006) | 4.74e-17* | 0.003 (0.006) | 0.607 |
| Superior longitudinal fasciculus | FA | -0.017 (0.007) | 0.013* | -0.013 (0.007) | 0.047 | -0.017 (0.007) | 0.014 |
|  | MD | 0.034 (0.007) | 5.70e-07* | 0.022 (0.007) | 6.86e-04* | 0.008 (0.007) | 0.25 |
| Superior thalamic radiation | FA | 0.001 (0.007) | 0.856 | -0.012 (0.007) | 0.093 | -0.009 (0.007) | 0.178 |
|  | MD | 0.038 (0.006) | 4.31e-10* | 0.039 (0.006) | 1.98e-11* | 0.007 (0.006) | 0.24 |
| Uncinate fasciculus | FA | -0.016 (0.006) | 0.014* | -6.78e-05 (0.006) | 0.991 | -0.015 (0.006) | 0.02 |
|  | MD | 0.026 (0.006) | 1.90e-05* | 0.01 (0.006) | 0.098 | 0.007 (0.006) | 0.257 |
| Forceps major | FA | -0.003 (0.007) | 0.719 | -0.013 (0.007) | 0.06 | -0.025 (0.007) | 3.44e-04* |
|  | MD | 0.017 (0.007) | 0.017* | 0.011 (0.007) | 0.105 | 0.015 (0.007) | 0.031 |
| Forceps minor | FA | -0.027 (0.007) | 1.40e-04* | -0.015 (0.007) | 0.027 | -0.009 (0.007) | 0.216 |
|  | MD | 0.031 (0.007) | 5.39e-06* | 0.023 (0.007) | 5.83e-04* | 0.006 (0.007) | 0.336 |
| Middle cerebellar peduncle | FA | 0.026 (0.007) | 5.25e-04* | 0.015 (0.007) | 0.034 | -0.005 (0.007) | 0.465 |
|  | MD | 0.051 (0.007) | 1.01e-12* | 0.059 (0.007) | 2.88e-17* | 0.001 (0.007) | 0.88 |

Table S6. The effects of stressful life events (SLE), neuroticism, smoking status, and alcohol consumption as covariates for NODDI measures. The test was conducted on the full sample of IDP from UK Biobank imaging team, without outlier exclusion or phenotypic data merging Betas were standardised effect sizes. P values were un-corrected p values. All p_corrected_<0.05 were marked by asterixis. FDR correction was applied on all the tests for a dMRI measure.

|  |  | **Alcohol consumption** | | **Smoking** | | **SLE** | |
| --- | --- | --- | --- | --- | --- | --- | --- |
| **Tract name** | **Measure** | **Beta (Std)** | **p** | **Beta (Std)** | **p** | **Beta (Std)** | **p** |
| g.Total | ICVF | -0.032 (0.007) | 6.28e-06* | -0.023 (0.007) | 7.70e-04* | -0.009 (0.007) | 0.205 |
|  | ISOVF | 0.036 (0.007) | 8.40e-08* | 0.023 (0.007) | 3.80e-04* | 0.004 (0.007) | 0.573 |
|  | OD | -0.029 (0.007) | 2.65e-05* | -0.009 (0.007) | 0.164 | 0.008 (0.007) | 0.228 |
| g.AF | ICVF | -0.034 (0.007) | 2.40e-06* | -0.022 (0.007) | 0.001* | -0.009 (0.007) | 0.214 |
|  | ISOVF | 0.012 (0.007) | 0.092 | 0.009 (0.007) | 0.178 | 0.006 (0.007) | 0.39 |
|  | OD | -0.014 (0.007) | 0.041* | 0.003 (0.006) | 0.675 | 0.009 (0.007) | 0.176 |
| g.TR | ICVF | -0.037 (0.007) | 1.74e-07* | -0.031 (0.007) | 4.50e-06* | -0.009 (0.007) | 0.212 |
|  | ISOVF | 0.052 (0.007) | 9.28e-16* | 0.034 (0.006) | 5.44e-08* | 0.003 (0.006) | 0.641 |
|  | OD | -0.03 (0.007) | 6.89e-06* | -0.013 (0.007) | 0.052 | 0.001 (0.007) | 0.855 |
| g.PF | ICVF | -0.01 (0.007) | 0.171 | -0.007 (0.007) | 0.283 | -0.007 (0.007) | 0.296 |
|  | ISOVF | 0.043 (0.007) | 1.54e-09* | 0.044 (0.007) | 2.30e-10* | -0.004 (0.007) | 0.554 |
|  | OD | -0.049 (0.007) | 3.97e-12* | -0.041 (0.007) | 3.40e-09* | -0.001 (0.007) | 0.875 |
| Acoustic radiation | ICVF | -0.024 (0.007) | 6.10e-04* | -0.012 (0.007) | 0.08 | -0.009 (0.007) | 0.162 |
|  | ISOVF | -0.012 (0.006) | 0.046 | -0.022 (0.006) | 1.93e-04* | 0.002 (0.006) | 0.715 |
|  | OD | -0.02 (0.006) | 0.001* | -0.029 (0.006) | 8.22e-07* | -0.002 (0.006) | 0.739 |
| Anterior thalamic radiation | ICVF | -0.028 (0.007) | 2.92e-05* | -0.011 (0.007) | 0.09 | -0.007 (0.007) | 0.256 |
|  | ISOVF | 0.029 (0.006) | 6.19e-06* | 0.029 (0.006) | 2.87e-06* | 0.005 (0.006) | 0.393 |
|  | OD | 3.79e-04 (0.007) | 0.955 | 0.002 (0.007) | 0.778 | 0.002 (0.007) | 0.743 |
| Cingulate gyrus part of cingulum | ICVF | -0.027 (0.007) | 9.78e-05* | -0.02 (0.007) | 0.002* | -0.009 (0.007) | 0.197 |
|  | ISOVF | -0.008 (0.006) | 0.206 | 2.94e-06 (0.006) | 1 | 0.002 (0.006) | 0.701 |
|  | OD | 0.014 (0.006) | 0.021* | 0.015 (0.006) | 0.011* | 2.07e-04 (0.006) | 0.972 |
| Parahippocampal part of cingulum | ICVF | -0.007 (0.007) | 0.318 | 9.61e-04 (0.006) | 0.882 | 0.004 (0.006) | 0.567 |
|  | ISOVF | 0.007 (0.006) | 0.238 | 0.008 (0.006) | 0.17 | 0.004 (0.006) | 0.504 |
|  | OD | -0.011 (0.006) | 0.072 | 0.004 (0.006) | 0.524 | 0.006 (0.006) | 0.283 |
| Corticospinal tract | ICVF | -0.013 (0.007) | 0.081 | -0.017 (0.007) | 0.015* | -0.01 (0.007) | 0.151 |
|  | ISOVF | 0.011 (0.007) | 0.113 | -0.006 (0.006) | 0.373 | 0.001 (0.006) | 0.832 |
|  | OD | -0.044 (0.007) | 3.52e-11* | -0.026 (0.006) | 5.85e-05* | -0.005 (0.006) | 0.421 |
| Inferior fronto occipital fasciculus | ICVF | -0.031 (0.007) | 8.86e-06* | -0.022 (0.007) | 0.001* | -0.008 (0.007) | 0.264 |
|  | ISOVF | 0.019 (0.007) | 0.006* | -0.003 (0.007) | 0.704 | 0.001 (0.007) | 0.848 |
|  | OD | -0.029 (0.007) | 1.55e-05* | -0.015 (0.006) | 0.019* | 0.002 (0.006) | 0.811 |
| Inferior longitudinal fasciculus | ICVF | -0.03 (0.007) | 2.28e-05* | -0.024 (0.007) | 3.95e-04* | -0.007 (0.007) | 0.329 |
|  | ISOVF | 0.029 (0.007) | 2.17e-05* | -0.001 (0.007) | 0.867 | -2.40e-05 (0.007) | 0.997 |
|  | OD | -0.018 (0.007) | 0.005* | 0.003 (0.006) | 0.582 | 0.005 (0.006) | 0.442 |
| Medial lemniscus | ICVF | 0.021 (0.006) | 5.27e-04* | 0.01 (0.006) | 0.104 | 0.004 (0.006) | 0.452 |
|  | ISOVF | 0.013 (0.006) | 0.046 | 0.004 (0.006) | 0.526 | -0.008 (0.006) | 0.163 |
|  | OD | 0.009 (0.007) | 0.207 | 0.019 (0.007) | 0.003* | 0.008 (0.007) | 0.239 |
| Posterior thalamic radiation | ICVF | -0.035 (0.007) | 6.42e-07* | -0.038 (0.007) | 1.34e-08* | -0.007 (0.007) | 0.315 |
|  | ISOVF | 0.051 (0.006) | 1.41e-15* | 0.03 (0.006) | 1.50e-06* | 0.001 (0.006) | 0.838 |
|  | OD | -0.013 (0.006) | 0.029* | 0.009 (0.006) | 0.115 | 0.01 (0.006) | 0.097 |
| Superior longitudinal fasciculus | ICVF | -0.034 (0.007) | 2.93e-06* | -0.022 (0.007) | 0.001* | -0.009 (0.007) | 0.174 |
|  | ISOVF | 0.018 (0.007) | 0.008* | 0.007 (0.006) | 0.269 | 0.004 (0.006) | 0.546 |
|  | OD | -0.027 (0.006) | 1.27e-05* | -0.013 (0.006) | 0.027* | 0.009 (0.006) | 0.143 |
| Superior thalamic radiation | ICVF | -0.039 (0.007) | 2.27e-08* | -0.038 (0.007) | 1.80e-08* | -0.009 (0.007) | 0.162 |
|  | ISOVF | 0.02 (0.006) | 5.87e-04* | 0.017 (0.006) | 0.002* | 0.004 (0.006) | 0.455 |
|  | OD | -0.034 (0.007) | 4.04e-07* | -0.016 (0.006) | 0.01* | -7.12e-04 (0.006) | 0.912 |
| Uncinate fasciculus | ICVF | -0.033 (0.007) | 1.17e-06* | -0.016 (0.006) | 0.014* | -0.01 (0.006) | 0.13 |
|  | ISOVF | 0.006 (0.007) | 0.391 | -0.003 (0.006) | 0.579 | 0.007 (0.006) | 0.294 |
|  | OD | -0.002 (0.006) | 0.798 | -0.002 (0.006) | 0.796 | 0.011 (0.006) | 0.068 |
| Forceps major | ICVF | -0.03 (0.007) | 4.18e-05* | -0.024 (0.007) | 6.38e-04* | -0.014 (0.007) | 0.052 |
|  | ISOVF | 0.003 (0.007) | 0.724 | -0.003 (0.007) | 0.704 | 0.013 (0.007) | 0.055 |
|  | OD | -0.034 (0.007) | 2.49e-06* | -0.019 (0.007) | 0.006* | 0.017 (0.007) | 0.014 |
| Forceps minor | ICVF | -0.035 (0.007) | 8.30e-07* | -0.017 (0.007) | 0.015* | -0.005 (0.007) | 0.495 |
|  | ISOVF | 0.003 (0.007) | 0.676 | 0.014 (0.007) | 0.042 | 0.008 (0.007) | 0.254 |
|  | OD | -0.014 (0.007) | 0.052 | -0.002 (0.007) | 0.751 | -6.06e-04 (0.007) | 0.932 |
| Middle cerebellar peduncle | ICVF | -6.47e-04 (0.007) | 0.926 | 0.004 (0.007) | 0.553 | -0.004 (0.007) | 0.603 |
|  | ISOVF | 0.047 (0.007) | 1.42e-10* | 0.057 (0.007) | 4.16e-16* | 0.002 (0.007) | 0.826 |
|  | OD | -0.037 (0.007) | 3.22e-07* | -0.033 (0.007) | 2.30e-06* | 4.28e-04 (0.007) | 0.951 |

Table S7. Comparing models with and without the cross-sectional measure. H0 model: Imaging variables~covariates+Mean+Variability, and H1 model: Imaging variables~covariates+Mean+Variability+Cross-sectional measure. ANOVA was utilised to test if adding the cross-sectional measure as an independent variable gives significantly larger variance explained by the model compared to the H0 model. Dependent variables were MD in the tract categories/tracts that were found associated with the cross-sectional measure. Significant p values are highlighted in red.

| **Method** | **Model** | **Independent variable(s)** | **Dependent variable** | **RSS** | **F test Df** | **F** | **p** |
| --- | --- | --- | --- | --- | --- | --- | --- |
| glm | H0 | Mean+Variability | g.MD.Total | 11707.16 | -- | -- | -- |
|  | H1 | Mean+Variability+Cross-sectional | g.MD.Total | 11700.27 | 1 | 8.659143 | 0.003259 |
|  | H0 | Mean+Variability | g.MD.AF | 12311.36 | -- | -- | -- |
|  | H1 | Mean+Variability+Cross-sectional | g.MD.AF | 12304.74 | 1 | 7.902749 | 0.004943 |
|  | H0 | Mean+Variability | g.MD.TR | 10495.86 | -- | -- | -- |
|  | H1 | Mean+Variability+Cross-sectional | g.MD.TR | 10491.75 | 1 | 5.745006 | 0.016548 |
|  | H0 | Mean+Variability | g.MD.PF | 13164.31 | -- | -- | -- |
|  | H1 | Mean+Variability+Cross-sectional | g.MD.PF | 13162.36 | 1 | 2.180021 | 0.139834 |
|  | H0 | Mean+Variability | Middle Cerebellar Peduncle | 13402.99 | -- | -- | -- |
|  | H1 | Mean+Variability+Cross-sectional | Middle Cerebellar Peduncle | 13402.25 | 1 | 0.820329 | 0.365099 |
|  |  |  |  | **Deviance** | **Chi test Df** | **Chisq** | **p** |
| lme | H0 | Mean+Variability | Anterior Thalamic Radiation | 53866.2 | -- | -- | -- |
|  | H1 | Mean+Variability+Cross-sectional | Anterior Thalamic Radiation | 53864.45 | 1 | 1.747914 | 0.186139 |
|  | H0 | Mean+Variability | Uncinate Fasciculus | 65881.34 | -- | -- | -- |
|  | H1 | Mean+Variability+Cross-sectional | Uncinate Fasciculus | 65878.74 | 1 | 2.600238 | 0.106848 |
|  | H0 | Mean+Variability | Superior Longitudinal Fasciculus | 60468.18 | -- | -- | -- |
|  | H1 | Mean+Variability+Cross-sectional | Superior Longitudinal Fasciculus | 60462.46 | 1 | 5.726051 | 0.016715 |
|  | H0 | Mean+Variability | Inferior Fronto Occipital Fasciculus | 64253.47 | -- | -- | -- |
|  | H1 | Mean+Variability+Cross-sectional | Inferior Fronto Occipital Fasciculus | 64249.67 | 1 | 3.798664 | 0.051293 |
|  | H0 | Mean+Variability | Superior Thalamic Radiation | 56451.53 | -- | -- | -- |
|  | H1 | Mean+Variability+Cross-sectional | Superior Thalamic Radiation | 56444.53 | 1 | 6.99626 | 0.008168 |
|  | H0 | Mean+Variability | Inferior Longitudinal Fasciculus | 61972.76 | -- | -- | -- |
|  | H1 | Mean+Variability+Cross-sectional | Inferior Longitudinal Fasciculus | 61966.72 | 1 | 6.037542 | 0.014005 |
|  | H0 | Mean+Variability | Corticospinal Tract | 71746.14 | -- | -- | -- |
|  | H1 | Mean+Variability+Cross-sectional | Corticospinal Tract | 71737.03 | 1 | 9.117795 | 0.002531 |
|  | H0 | Mean+Variability | Acoustic Radiation | 79033.73 | -- | -- | -- |
|  | H1 | Mean+Variability+Cross-sectional | Acoustic Radiation | 79027.2 | 1 | 6.535737 | 0.010573 |
|  | H0 | Mean+Variability | Cingulate Gyrus Part Of Cingulum | 64031.44 | -- | -- | -- |
|  | H1 | Mean+Variability+Cross-sectional | Cingulate Gyrus Part Of Cingulum | 64022.83 | 1 | 8.611179 | 0.003341 |

Table S8. Comparing models with and without the longitudinal measures. H0 model: Imaging variables~covariates+Cross-sectional measure, and H1 model: Imaging variables~covariates+Mean+Variability+Cross-sectional measure. ANOVA was utilised to test if adding the cross-sectional measure as an independent variable gives significantly larger variance explained by the model compared to the H0 model. Dependent variables were MD in the tract categories/tracts that were found associated with the cross-sectional measure. Significant p values are highlighted in red.

| **Method** | **Model** | **Independent variable(s)** | **Dependent variable** | **RSS** | **F test Df** | **F** | **p** |
| --- | --- | --- | --- | --- | --- | --- | --- |
| glm | H0 | Cross-sectional | g.MD.Total | 11701.79 | -- | -- | -- |
|  | H1 | Mean+Variability+Cross-sectional | g.MD.Total | 11700.27 | 2 | 0.959116 | 0.383256 |
|  | H0 | Cross-sectional | g.MD.AF | 12306.15 | -- | -- | -- |
|  | H1 | Mean+Variability+Cross-sectional | g.MD.AF | 12304.74 | 2 | 0.840598 | 0.431473 |
|  | H0 | Cross-sectional | g.MD.TR | 10493.18 | -- | -- | -- |
|  | H1 | Mean+Variability+Cross-sectional | g.MD.TR | 10491.75 | 2 | 0.997852 | 0.368695 |
|  | H0 | Cross-sectional | g.MD.PF | 13172.09 | -- | -- | -- |
|  | H1 | Mean+Variability+Cross-sectional | g.MD.PF | 13162.36 | 2 | 5.430353 | 0.00439 |
|  | H0 | Cross-sectional | Middle Cerebellar Peduncle | 13410.69 | -- | -- | -- |
|  | H1 | Mean+Variability+Cross-sectional | Middle Cerebellar Peduncle | 13402.25 | 2 | 4.625691 | 0.009811 |
|  |  |  |  | **Deviance** | **Chi Df** | **Chisq** | **Pr(>Chisq)** |
| lme | H0 | Cross-sectional | Anterior Thalamic Radiation | 53866.22 | -- | -- | -- |
|  | H1 | Mean+Variability+Cross-sectional | Anterior Thalamic Radiation | 53864.45 | 2 | 1.776644 | 0.411345 |
|  | H0 | Cross-sectional | Uncinate Fasciculus | 65878.81 | -- | -- | -- |
|  | H1 | Mean+Variability+Cross-sectional | Uncinate Fasciculus | 65878.74 | 2 | 0.070895 | 0.965174 |
|  | H0 | Cross-sectional | Superior Longitudinal Fasciculus | 60463.21 | -- | -- | -- |
|  | H1 | Mean+Variability+Cross-sectional | Superior Longitudinal Fasciculus | 60462.46 | 2 | 0.757896 | 0.684581 |
|  | H0 | Cross-sectional | Inferior Fronto Occipital Fasciculus | 64250.01 | -- | -- | -- |
|  | H1 | Mean+Variability+Cross-sectional | Inferior Fronto Occipital Fasciculus | 64249.67 | 2 | 0.34637 | 0.840982 |
|  | H0 | Cross-sectional | Superior Thalamic Radiation | 56447.14 | -- | -- | -- |
|  | H1 | Mean+Variability+Cross-sectional | Superior Thalamic Radiation | 56444.53 | 2 | 2.609169 | 0.271285 |
|  | H0 | Cross-sectional | Inferior Longitudinal Fasciculus | 61968.44 | -- | -- | -- |
|  | H1 | Mean+Variability+Cross-sectional | Inferior Longitudinal Fasciculus | 61966.72 | 2 | 1.718461 | 0.423488 |
|  | H0 | Cross-sectional | Corticospinal Tract | 71740.46 | -- | -- | -- |
|  | H1 | Mean+Variability+Cross-sectional | Corticospinal Tract | 71737.03 | 2 | 3.428328 | 0.180114 |
|  | H0 | Cross-sectional | Acoustic Radiation | 79029.9 | -- | -- | -- |
|  | H1 | Mean+Variability+Cross-sectional | Acoustic Radiation | 79027.2 | 2 | 2.705615 | 0.258514 |
|  | H0 | Cross-sectional | Cingulate Gyrus Part Of Cingulum | 64025.77 | -- | -- | -- |
|  | H1 | Mean+Variability+Cross-sectional | Cingulate Gyrus Part Of Cingulum | 64022.83 | 2 | 2.939637 | 0.229967 |

**References**

1. Rosseel Y (2012): lavaan : an R package for structural equation modeling and more Version 0 . 5-12 ( BETA ). *J Stat Softw*. 48: 1–36.

2. Clarke T-K, Adams MJ, Davies G, Howard DM, Hall LS, Padmanabhan S, *et al.* (2017): Genome-wide association study of alcohol consumption and genetic overlap with other health-related traits in UK Biobank (N=112 117). *Mol Psychiatry*. 22: 1376–1384.

3. Smith SM, Nichols TE (2018): Statistical Challenges in “Big Data” Human Neuroimaging NeuroView. *Neuron*. 97: 263–268.
